# Microengineered 2D and 3D modular neuronal networks represent structure-function relationship

**DOI:** 10.1101/2023.04.07.535751

**Authors:** Rouhollah Habibey, Johannes Striebel, Roshanak Latiftikhereshki, Felix Schmieder, Shahrzad Latifi

**Affiliations:** Universitäts-Augenklinik Bonn, University of Bonn, Dep. of Ophthalmology, Ernst-Abbe-Straße 2, D-53127 Bonn, Germany; CRTD - Center for Regenerative Therapies TU Dresden, 01307 Dresden, Germany; Department of Computer Engineering, Faculty of Engineering, Kermanshah Branch, Azad University, Kermanshah, Iran; Laboratory of Measurement and Sensor System Technique, Faculty of Electrical and Computer Engineering, TU Dresden, Helmholtzstraße 18, 01069 Dresden, Germany; Department of Neurology, David Geffen School of Medicine, University of California Los Angeles, Los Angeles, California, USA; Department of Neuroscience, Rockefeller Neuroscience Institute West Virginia University, Morgantown, WV, 26506, USA

**Keywords:** Microengineering, Microphysiological systems, Microfluidics, Modular networks, Structure-function relationship, 3D neuronal culture.

## Abstract

Brain function is substantially linked to the highly organized structure of neuronal networks. Emerging three-dimensional (3D) neuronal cell culture technologies attempt to mimic the complexity of brain circuits as *in vitro* microphysiological systems. Nevertheless, structures of *in vitro* assembled neuronal circuits often varies between samples and changes over time that makes it challenging to reliably record network functional output and link it to the network structure. Hence, engineering neuronal structures with pre- defined geometry and reproducible functional features are essential to model *in vivo* neuronal circuits in a robust way. Here, we engineered thin microchannel devices to assemble 2D and 3D modular networks. Microchannel devices were coupled with multi-electrode array (MEA) electrophysiology system to enable long-term electrophysiology recordings from microengineered circuits. Each network was composed of 64 micromodules which were connected through micron size channels to their adjacent modules. Microstructures physically confined neurons to the recording electrodes that considerably enhanced the electrophysiology readout efficiency. In addition, microstructures preserved modular network structure over weeks. Modular circuits within microfluidic devices showed consistent spatial patterns of activity over weeks, which was missing in the randomly formed circuits. Number of physical connections per module was shown to be influencing the measured activity and functional connectivity parameters, that represents the impact of network structure on its functional output. We show that microengineered 3D modular networks with a profound activity and higher number of functional connections recapitulate key functional features of developing cortex. Structurally and functionally stable 2D and 3D network mimic the modular architecture of brain circuits and offers a robust and reproducible *in vitro* microphysiolopgical system to serve basic and translational neuroscience research.

## Introduction

Sophisticated cognitive features of the human brain are derived from an enormous number of neurons and their intricate but organized connections [1]. Mapping the network structural connectivity attempts to resolve delicate wiring diagrams and network architecture at synapse levels [2,3]. These data remarkably support the emerging field of network neuroscience to link the organizational attributes of the brain circuits with their functional outputs [2,4,5]. Functional connectivity maps are based on statistical measures [5] that identify correlated activities between different regions of the network [4]. However, functional connectivity maps of *in vivo* neuronal networks are highly dynamic and often subjected to external stimuli or modulated by the presence of complex autocrine and paracrine signaling cues [4,6,7]. Therefore, despite a significant correlation between structure and function [8], often it is challenging to estimate network function directly from its structure in the *in vivo* setups [7,8].

From a functional standpoint, bottom-up neuronal engineering endeavors to generate *in vitro* neuronal circuits with reproducible activity and connectivity patterns, resembling *in vivo* networks [9–11]. Recent advances in engineering neuronal networks with reduced complexity provide applicable control over network structure that can be exploited to study the structure-function relationship [11–14]. Position of neurons, their growth direction, and connectivity can be controlled by patterning the substrate with cell adhesive proteins or physically confining neurons and neurites in microfluidic or microchannel devices [9,11,15,16]. Engineered neuronal circuits with a pre-defined structure are integrated with microscopy and electrophysiology readout systems to monitor network morphology and function in parallel [9,13,14,17– 21].

The *in vitro* network function is mainly studied multi-electrode arrays (MEAs) electrophysiology and optical tools like calcium imaging and optogenetics [11,14,15,22,23]. Non-invasive *in vitro* electrophysiology tools for multi-site long-term recordings, including transparent MEAs and high-density MEAs, are commercially available [13,22,24–27]. They have been applied for mapping the functional connectivity patterns in randomly connected 2D neuronal networks [14,26,28,29]. However, random networks do not recapitulate the modular structure of the brain circuits [28]. In addition, *in vitro* network structure undertakes changes by culture age due to the shifts in the position of the neuronal cell body and axonal branches [14,21,27]. It means that a specific electrode does not necessarily record from same neurons or part of the circuit over weeks and months of network development [27]. This can lead to inconsistency in tracking activity and functional features, particularly for long-term experiments [27]. Therefore, engineered networks with stable structural motifs confined to MEA electrodes can improve the reliability of functional data [30].

Two-dimensional networks with modular structures and diverse connectivity configurations can be assembled on MEA substrate [14,15,31,32]. For instance, microcontact printing (µCP) has been applied to localize neuronal cell bodies to recording electrodes and define their synaptic communication through connecting neurite-adhesive strips [31,33,34]. Diverse communication geometries between nodes, the population of neurons on electrodes, were generated by changes in the design of the stamps [35,36]. Yet, surface-bonded cell adhesive patterns often degrade leading to changes in network morphology on the long run [29,36–38]. Engineered cell-repellent hydrophobic regions are also prone to be coated with cell-secreted materials that attract the unwanted growth of neurites outside of the patterned structures [11,29,37]. To overcome these issues, physical confinements can be applied to permanently hold neurons and their branches in desired configurations [16,39–41]. Polydimethylsiloxane (PDMS)-based microfluidic devices aligned on a glass substrate or coupled to the MEA chips provide physical support for neuronal network formation [16,30,32,39,42]. Integrated PDMS microchannels and low- or high-density MEA devices have served as a versatile platform to study long-term functional properties of developing neuronal networks [30,43,44], to generate a mechanically and chemically stable microenvironment [9,21,30], and to enhance the viability of the designed circuits with a limited number of neurons [11,45,46].

Conventional 2D neuronal culture systems have been accompanied by progress in 3D models to mimic the complex microarchitecture of brain circuitry *in vitro* [47,48]. 3D neuronal networks are either constructed by supporting scaffolds like hydrogel, colloids, and microfluidics or autonomously assembled as cell aggregates, spheroids, and organoids [11,18,49,50]. Microfluidics offer an improved microphysiological environment and precise control of culturing conditions, while organoids are superior in recapitulating the macroscale organization of the brain [11,48,51]. Functional measurements from 3D engineered networks have been mainly limited to flat MEAs or calcium imaging [52–54], apart from the recently developed 3D MEAs that can record network dynamics within 3D neuronal tissue [55]. To probe the activity dynamics of 3D networks, most groups have assembled a bulk 3D structure covering the MEA sensing areas and spaces between electrodes [53,56,57]. Regardless of their many advantages over 2D models, these 3D networks were made of randomly distributed and interconnected neurons [58,59]. Lately, some groups engineered modular 2D networks on MEAs by changing the degree of cellular aggregation or using PDMS microstructures to adjust the spatial arrangement of neurons [44,60]. These networks mimicked the modular microarchitecture of the brain circuits and exhibited improved functional resilience against chemical perturbation of excitatory synapses [60].

Here, we present a straightforward method to construct 2D and 3D modular networks with defined connectivity patterns within PDMS microchannel devices coupled to MEA electrodes. Modular networks were prepared with a significantly lower number of neurons compared to high-density random networks. However, modular 2D networks represented comparable activity, and modular 3D networks exhibited strong functional features resembling neonatal cortical networks. We show how the spatial arrangement of neurons in the engineered circuits enhanced the percentage of recordable units that are mostly inaccessible in recordings from random networks. In-depth analysis of our data revealed a relationship between predefined network structure and its function both in 2D and 3D modular networks. Most importantly, the stable structure of engineered circuits was reflected in consistent spatial patterns of activity as well as connectivity maps at different culture ages. Random networks on the other hand showed distinctive patterns of activity at each recording week. This approach can be extended for engineering neural networks derived from human stem cells to generate networks with reproducible activity and functional connectivity characteristics to model functional phenotype of neurological disorders on-chip.

## Materials and methods

### Fabrication of the PDMS microchannel device

The details of microchannel device fabrication have been discussed previously [30,61]. We applied a modified approach (Fig 1A). Briefly, SU-8 templates defining reverse geometries of microchannel devices were fabricated in two layers on a 4-inch silicon wafer (Si-Mat). To fabricate the template in two layers, SU- 8 5 and SU-8 50 (MicroChem) were subsequently spin-coated on the wafer at different heights (5 µm for microchannels and 100 µm for microwells; Ws-650Sz Spin Coater, Laurell Technologies). Photo-patterning on the two layers was done in a mask aligner (MJB4, SUSS MicroTec) through printed high-definition transparency masks (Repro S.r.l.). Stylus profiler (Wyko NT1100, Veeco) and quantitative microscopy (Leica DM IL LED Inverted, Leica Microsystems CMS GmbH) were used to determine the physical dimensions of the fabricated SU-8 master.

**Fig. 1.**
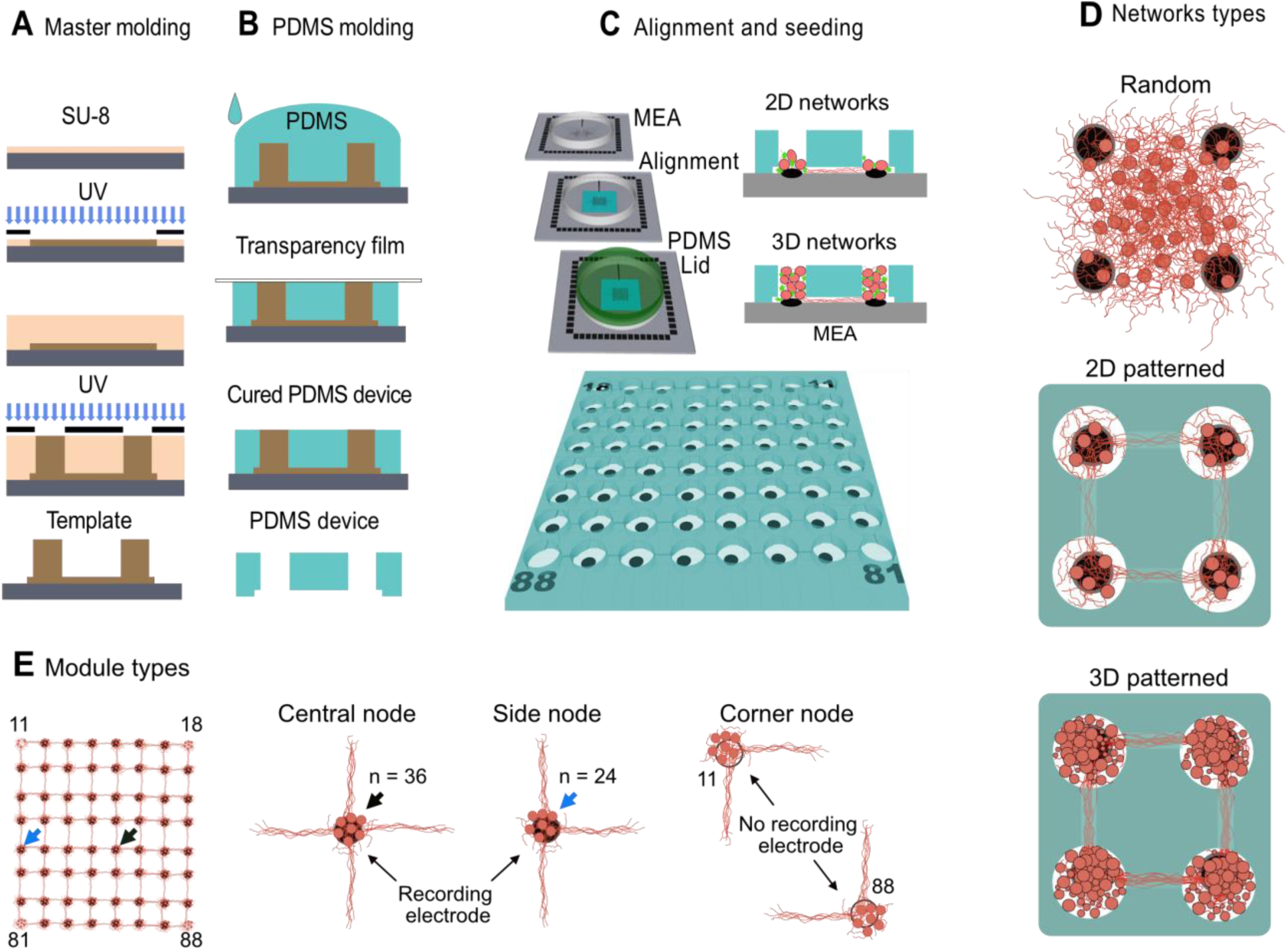
Steps to engineer 2D and 3D patterned neural networks. **A**) Photolithography and fabrication of the SU-8 template in 2 layers. First and second layers were fabricated with 5 µm and 100 µm heights, respectively. **B**) Fabrication of the PDMS devices on SU-8 template. Through holes were provided by leaving a transparency film on top of the bigger structures. **C**) PDMS device was aligned with MEA electrodes. After PDL coating, cells were seeded in microwells. **D**) Modular networks vs random network structure. 2D patterned networks seeded in low (2DL) small 2D modules are formed on top of each electrode composed of limited number of neurons and high (2DH) densities. 3D patterned networks formed by filling the whole microwell. **E**) Each 2D or 3D patterned culture was composed of 64 modules including 36 central nodes, 23 side nodes and 4 corner nodes. Each central node has 4 sides, each side node has 3 sides. Based on standard MEA configurations there is no electrode on the corner nodes.

PDMS pre-polymer and catalyst (Sylgard 184, Dow Corning) were mixed thoroughly (10:1), degassed in a desiccator to remove air bubbles, and poured on the original SU-8 template. After removing all air bubbles, a clean laser copier transparency film was positioned on top of the liquid PDMS (Fig 1B). A flat sponge and then a piece of glass were placed on the backside of the transparency film, then the sandwich of Su-8 template-PDMS-transparency film-sponge and glass was squeezed using Spring Clamp (Wolfcraft B3631 Fz60 60mm) to remove extra PDMS. This allowed the creation of through-holes that later served as PDMS microwells (Fig 1B and 1C). After curing PDMS at 80 °C for 30 minutes the replica was peeled off from the template and fully cured for 10 hours at 120 °C to remove all oligomer residues. The final device featured 64 microwells (Ø = 120 µm, h = 100 µm) that were connected by shallow microchannels (L = 80 µm, w = 30 µm and h = 5 µm; Fig 1B-C). These features were used to assemble modular 2D and 3D networks each including 64 modules (Fig 1D). Four modules in the corners and one module aligned to electrode 15, counter electrode, were not available for recording. All modules located in the edge of the design are connected through three microchannels to other modules and all central modules are connected to four neighboring modules (Fig 1D- E and Video 1).

### 2D and 3D neuronal cell culture in modular microchannel device

Embryos were harvested from pregnant Sprague Dawley rats (CD IGS, Charles River) at embryonic day 18 (E 18). After removing the meninges, the cortex was extracted and dissociated into single cells by incubating it for 10 minutes in 0.25% (w/v) trypsin in Hank’s Balanced Salt Solution (HBSS). After deactivation of trypsin, sequential trituration of the cell solution was applied to get a single cell suspension. The suspension was centrifuged (at 200 g for 5 minutes), the supernatant was removed, and cell pellets were re-suspended in complete Neurobasal medium (NBM) including B27 serum-free supplement (2%), penicillin/streptomycin (1 mM), and Glutamax (2 mM), all bought from Gibco/Life Technologies/Thermo Fisher Scientific.

Autoclaved (120 °C, 20 min) PDMS devices were either aligned with electrodes of standard MEA chips (30/200 ir, Multichannel Systems GmbH (MCS)) using a Stereo Microscope (Levenhuk 3ST Stereo Microscope) or placed on sterile cover glasses (Fig 1C and Video 1)[30]. Aligned devices and the surface of MEA chips or cover glasses were hydrophilized by oxygen plasma (2min, 60 W, 2.55 GHz, 0.3 mbar O2; Femto, Diener) before coating them with poly-D-lysine (PDL; 0.1 mg/ml). For the 2D group after adding 50 µl of PDL solution and removing the air bubbles, extra PDL was removed from the top of the device and microwells to limit coating to the MEA surface. In the 3D group, PDL solution was left inside the microwells to coat the walls of each microwell. PDL coating was incubated overnight and then washed with sterile MQ water three times. A day before seeding the cells, MQ water was removed and a PDMS mask with a large hole (Ø = 2mm) was aligned on top of the microchannel device (Video 1). The mask was later used to keep cell suspension on top of the device during the incubation period. The whole device was rinsed with complete NB media and air bubbles were removed. In the control group (N = 4 MEAs) with no PDMS device, after plasma treatment PDL 0.1 mg/ml was added (50 µl) to the center of the MEA, incubated overnight, and washed three times with MQ water. The same protocol of PDL coating was applied for coating the 18 mm coverslips in 12 well plates.

To create 2D low-density (2DL; N = 4 MEAs) and 2D high density (2DH; N = 4 MEAs) neuronal populations inside the microwells, 50 µl of 2000 cells/μl and 5000 cells/μl cell suspension was added, respectively. After 10 minutes of incubation, some cells were randomly but almost equally deposited into the microwells and the rest of them remained on the top of the device. Finally, the PDMS mask was gently removed and the cells on the top side of the PDMS device were washed out with cell culture media. To create 3D neuronal cultures in PDMS microchannel devices (N = 4 MEAs), 50 µl of neuronal cell suspension with 8,000 cells/μl was added and incubated for 10 minutes. The extra cell suspension was removed from the top of the device and the process of seeding was repeated two more times. This process stacked a mixture of neurons and astrocytes on top of each other inside individual microwells with the support of PDL-coated microwells. In the Control group, 50 µl neural cell suspension (2000 cells/μl; 100,000 cells per MEA) was added and incubated for 20 minutes. The same protocol of cell seeding was applied for cultures on 18 mm coverslips in 12 well plates (N=5 coverslip culture per group).

Serum-free cell culture medium was added to each MEA or coverslip in a 12-well plate. MEA rings were sealed with gas-permeable PDMS caps [62]. Cultures were always kept in an incubator except for sessions of electrophysiology recording, microscopy, and culture medium exchange. The medium was changed once a week by replacing 500 µl old media with fresh media. Cultures on coverslips were used for immunostaining and MEA cultures were used for electrophysiology recording. Bright-field microscopy images (20 images per culture) were prepared with high magnification (20x objectives) and were automatically stitched using ImageJ to visualize whole network morphology (Fig 2A and Supp Fig 1-4).

**Fig. 2.**
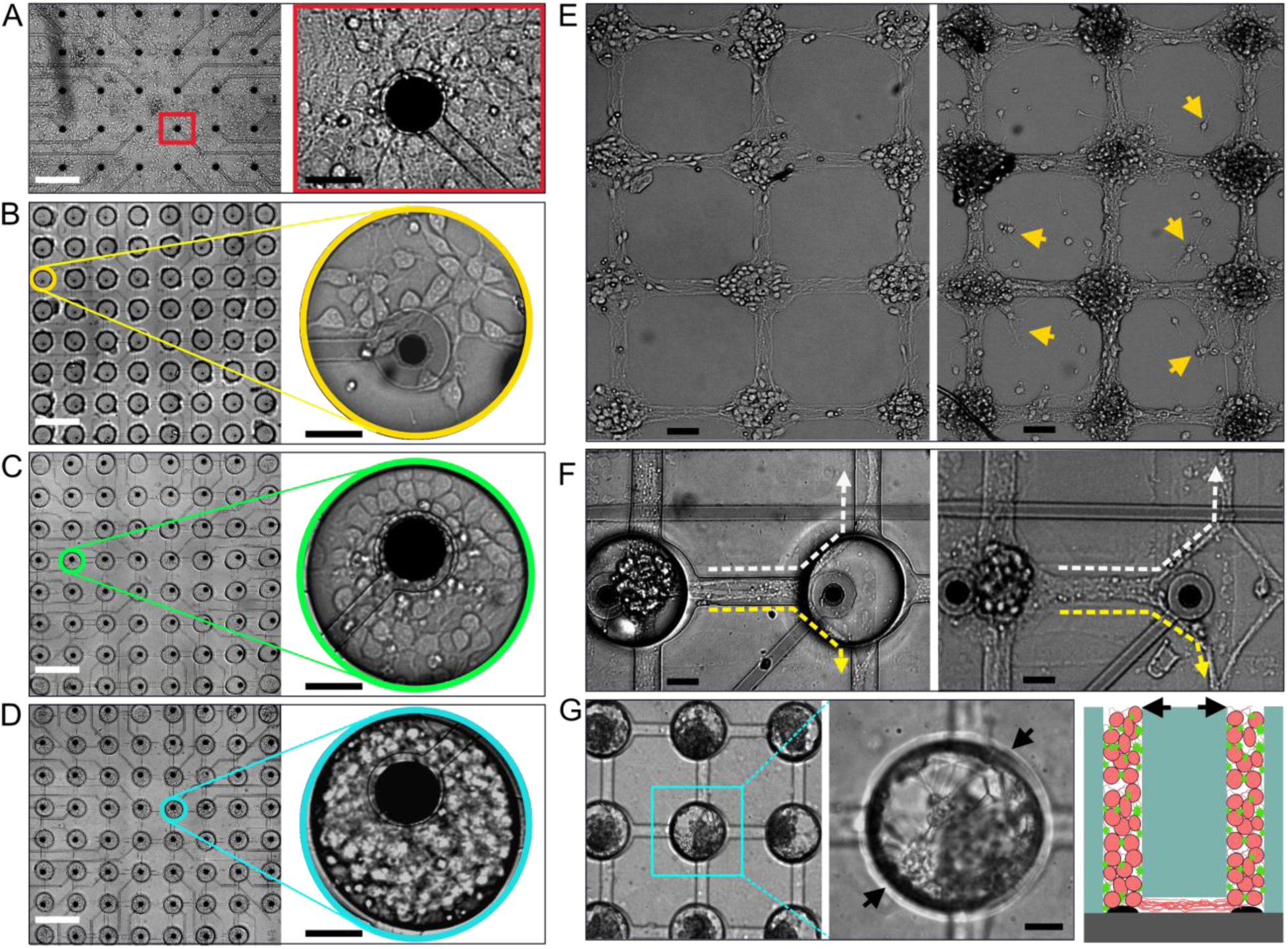
Bright-field microscopy of the network morphology. **A**) Bright-field microscopy image from a random network covering 30 electrodes and a magnified view of the network on top of a single electrode (red rectangle). **B**) Microscopy image from a low-density 2D patterned network (2DL); whole network and magnified view of a microwell (yellow circle). **C**) High-density 2D patterned network (2DH); whole network and magnified view of a network module (green circle). **D**) 3D patterned network and a selected 3D assembled neuronal structure inside a microwell (blue circle). **E**) Physical confinement by PDMS device is necessary for a stable network structure. Network structure just after removing device at 7 DIV (left) and 17 DIV (right) with neurons growing outside of the pre-defined structure (right; yellow arrows). **F**) Axonal branches take either a direct path (not shown here) or bend in the nodal point (arrows). Before (left) and after (right) removing PDMS device at 45 DIV. **G**) Image from the highest z-plane of the 3D network module at 35 DIV representing neuronal cell bodies and axonal branches attached to the microwell walls (black arrows). The magnified views of larger areas of each network type in all groups have been depicted in the supplementary File (Supp Fig 1-4)

### MEA electrophysiology

Standard 60-electrode MEA amplifier (MEA1060-upright, Multichannel systems) was used to capture electrical signals with a 25 kHz sampling frequency. During the recording, the culture media temperature was kept stable (37 °C) using a built-in thermal sensor and heating element that was controlled by an external temperature controller (HC-1, MCS). MC_Rack software user interface, provided by Multichannel Systems (MCS), was used for recording the raw data. Network activity was recorded at different days in vitro (DIV) starting from 10 DIV. At each DIV we left MEAs for 5 minutes on the MEA amplifier to adapt to the ambient conditions and then network activity was recorded for 10 minutes (Fig 1C). To reduce the effect of media exchange on recorded network activity, media was changed 3 days before recording [63]. Recorded signals were filtered offline in MC_Rack software by a second-order Bessel high-pass filter (cut- off at 100 Hz) and timestamps of detected action potentials (APs) were extracted by automatically adjusted negative threshold (−6.00 Standard deviation of the peak-to-peak noise). Extracted time stamps were imported in NeuroExplorer software (Nex Technologies) to calculate mean AP frequency on each electrode. Additionally, burst activity features and functional connectivity patterns were measured based on timestamp data.

### Immunofluorescence and confocal microscopy

Morphology of the engineered networks was visualized on 18 mm coverslips and selected MEA chips. Cultures were washed with warm PBS (phosphate buffered saline 1%), were fixed with 4% paraformaldehyde (PFA) plus 4% sucrose (10 min incubation time). Fixed samples were treated with 0.2% Triton X-100 (10 min incubation time). Later samples were incubated with blocking buffer including 2% goat serum (GS) and 3% bovine serum albumin (BSA), for 30–45 min. Subsequently, primary antibodies were applied and incubated for 60 minutes at RT, which was followed by incubation with secondary antibodies for 60 minutes in RT and dark conditions. Primary antibodies anti-rabbit β-tubulin III IgG (SAB4300623, Sigma) and monoclonal anti-glial fibrillary acidic protein (GFAP) IgG (G3893, Sigma) and secondary antibodies Alexa Fluor 647 and Alexa Fluor 488 were applied to label neurons and astrocytes, respectively. A mounting solution including DAPI was used to mark nuclei. Fluorescent images were prepared using either an upright fluorescence microscope (BX51, Olympus) or an inverted confocal microscope (Nikon A1+). Images were captured and stored using an Optronics Microfire microscope camera (2-megapixel, MBF). High-resolution images from large areas of the network on MEA or coverslip were generated by automatic stitching of mosaic images (20x) in ImageJ (Fig 3 and Supp Fig 5-7). To extract the three-dimensional structure of the patterned or random networks confocal images of all three wavelengths were taken in at least 33 z-planes (pitch = 3 µm), covering the whole thickness of the microchannel devices (100 µm). Z-stack images were assembled by ImageJ 3D viewer plugin to visualize the 3D structure of the network (Video 2 to Video 7) and to illustrate the z-axis profile of the constructed network (Fig 3C) [64].

**Fig. 3.**
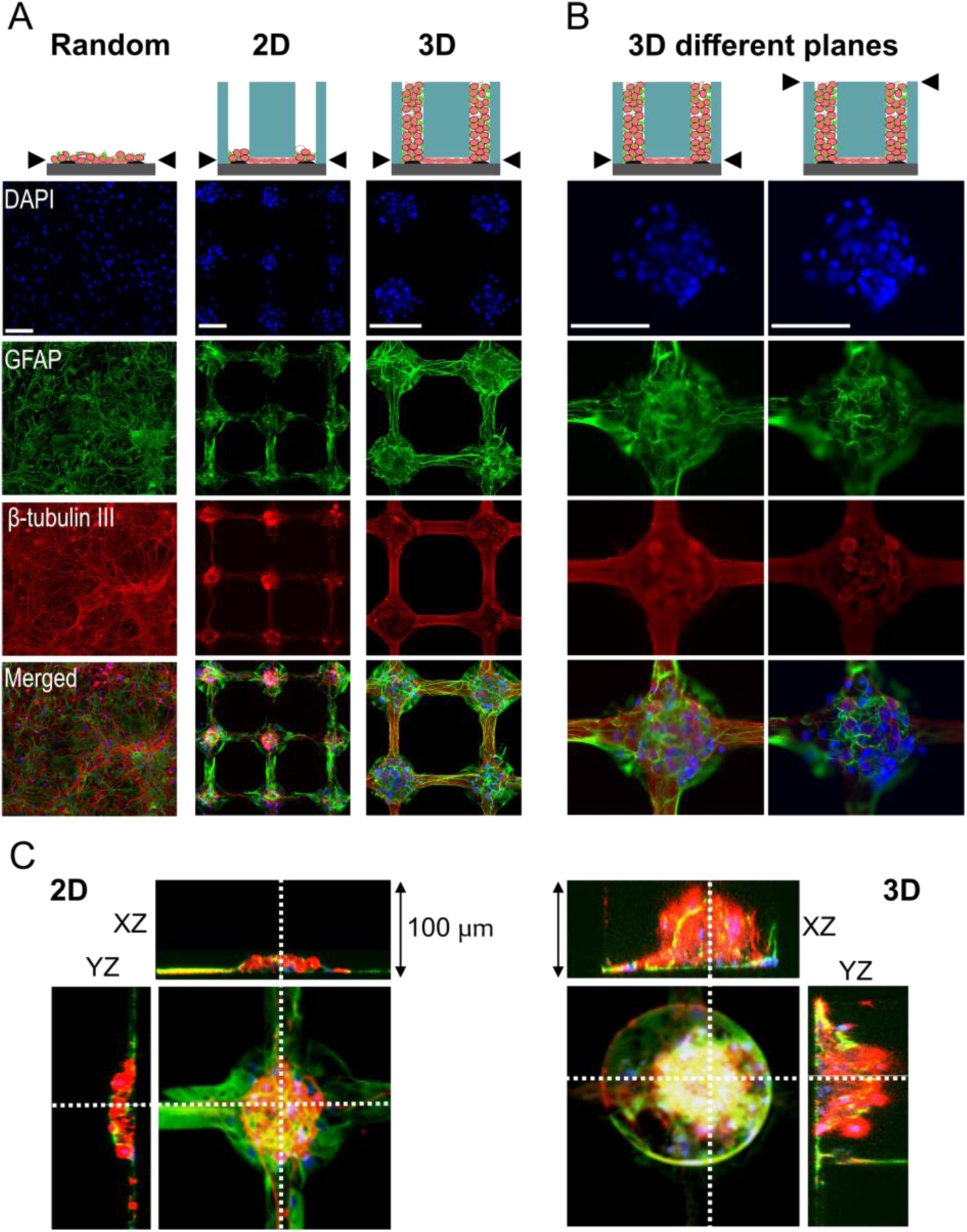
Immunofluorescence images of the network morphology. **A**) Random and 2D and 3D patterned networks stained for nuclei (DAPI), astrocytes (GFAP), and neurons (β-tubulin III). All images have been prepared from the microchannel plane (black arrows). **B**) Magnified view of the network in one microwell of a 3D culture in microchannel plane or 100 μm higher at the top surface of the device (black arrows). **C**) Orthogonal view of 2D patterned (left) and 3D patterned (right) networks from a selected microwell representing the thickness of the network in these two groups. Further details of the network structure in 2D and 3D patterned cultures on MEAs and coverslips have been depicted in the supplementary files (Supp Fig 2-7 and Video 2-7).

### Assessment of functional connection and connectivity maps

A cross-correlation based approach was used to estimate functional connectivity in each MEA culture. This approach has previously been validated for MEA recordings of *in vitro* neuronal populations [29,65]. Cross-correlation assesses the similarity of the spike train of one electrode (called target) relative to another electrode (called reference). The cross-correlation between each pair of active electrodes (x,y) was evaluated using the following cross-correlogram:

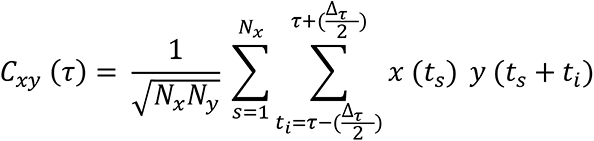

Where Nx and Ny represent the total number of spikes in x and y, ts is the duration of each spike in train x and Δτ the time window in which spikes in train y are detected. Here, a time window of 0.5 ms was applied.

A threshold was applied to the cross-correlations to retain only statistically significant links and to remove physiologically implausible values. A hard threshold defined as T=μ+*n*·σ was used, where μ and σ were the mean and the standard deviation computed among all the cross-correlation values, and *n* was an integer [7,14,28,66]. The output of thresholding step was a series of connectivity matrices where highest values (cross-correlation) were corresponding to the strongest connections between pairs of electrodes. A functional graph was obtained from these matrices for each MEA, which was weighted by cross-correlation values. For those electrode pairs that possessed the cross-correlation values below the threshold, no connection was drawn. These graphs show the physical distribution of the network on the MEAs which is corresponding to the distinct functional connections of electrodes.

### Statistical analysis

APs were detected by MC_Rack software and extracted time stamps were imported into NeuroExplorer (Nex Technologies) to calculate the AP frequency and different features of burst activity at each electrode. Electrodes not showing activity in particular days were set to zero to probe the development of activity features over time in different DIVs. Complex flares of recorded activity in each electrode were considered as burst activity if it met the following criteria: 20 ms maximum inter-spike interval to start the burst, 10 ms maximum inter-spike interval to end the burst, 10 ms minimum inter-burst interval, 20 ms minimum burst-duration with at least 4 spikes in each burst [21,30,62]. To extract unit data recorded by each electrode we used raw AP wavelets that have been detected and saved (threshold = −6 Standard deviation of the peak-to-peak noise, pre-trigger time = 1 ms, post-trigger time = 2 ms, and dead time = 2 ms). AP wavelets were imported into Plexon offline sorter (version 3.3.0) and were sorted with k-mean scan sorting. Based on visual inspection k-mean scan was more reliable than other algorithms on separating the neural units. Extracted units were exported again into NeuroExplorer for measuring the AP frequency in neural units at different groups.

Activity features including AP and burst frequency, burst duration, and percentage of APs in a burst were calculated in each electrode per DIV, averaged across all electrodes of the same group and compared with the control random networks, using mixed-effects analysis followed by Tukey’s multiple comparison test (Fig 4). The mean number of units per electrode (N= 4 MEA per group and n=236 electrodes per group) was calculated between 15 DIV and 45 DIV data and compared between groups based on Kruskal–Wallis one-way analysis of variance followed by Dunn’s multiple comparison test (Fig 4D). The average AP frequency of neural units detected by each electrode was calculated for 15 DIV to 45 DIV recordings and compared between groups based on Kruskal–Wallis one-way analysis of variance followed by Dunn’s multiple comparison test (Fig 4D). The percentage of neurons in each network available for recording was measured by the total number of neurons in each network and the total number of detected units in each MEA. Both values were divided by the number of electrodes per MEA (n=59), then the percentage of available units per electrode was calculated in each DIV. Lastly, the mean value of the percentage of available units per electrode between 15 DIV and 45 DIV was measured and compared between groups (n=236 electrodes per group; Fig 4D).

**Fig. 4.**
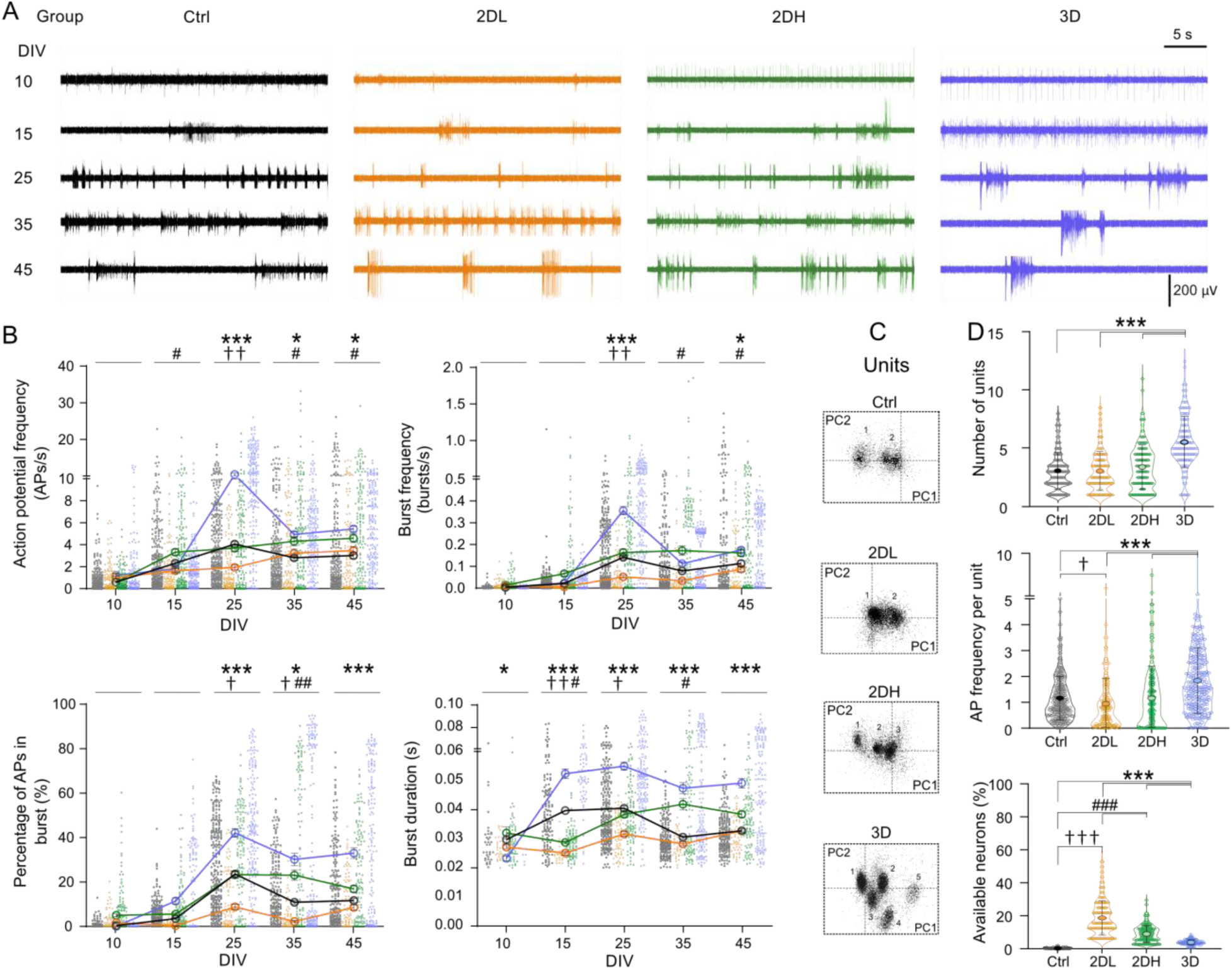
Activity profile in engineered 2D and 3D networks. **A**) Exemplar trace of activity in a selected electrode at different DIV in each group. **B**) Action potential (AP) and burst frequency, percentage of APs in the burst, and burst duration for all four conditions over time. Ctrl is represented in gray, 2DL in orange, 2DH in green and 3D in blue. Each point represents the average of the corresponding variable in one electrode a day. Data has been presented as Mean ± S.E.M (N= 4 MEAs, and n=236 electrodes per group). Data between groups were compared using mixed- effects analysis followed by Tukey’s multiple comparison test. 2DL *vs.* Ctrl († p < 0.05, †† p < 0.01), 2DH *vs.* Ctrl (# p < 0.05, ## p < 0.01), and 3D *vs.* Ctrl (* p < 0.05, ** p < 0.01, *** p < 0.001). **C**) Example neural units extracted based on k-means clustering in a selected electrode from each group at DIV 45. **D**) Number of detected units per electrode, AP frequency per unit, and percentage of neurons available for recording per electrode. Data in each group has been pooled from 15 DIV to 45 DIV. Data between groups were compared using Kruskal–Wallis one-way analysis of variance followed by Dunn’s multiple comparison test. 2DL *vs.* other groups († p < 0.05, ††† p < 0.001), 2DH *vs.* other groups (### p < 0.01), and 3D *vs.* other groups *** p < 0.001).

Extracted data from functional connectivity analysis including the number of connections between modules and the average strength of connections compared between groups (N= 4 MEA per group and n=236, n=118, n=177 and n=177 electrodes per group for Ctrl, 2DL, 2DH, and 3D groups, respectively) at different days was calculated using mixed-effects analysis followed by Tukey’s multiple comparison test (Fig 4)

To probe the spatial distribution of the network activity and its stability over time we tracked the trend of activity, including AP frequency and burst features, in individual electrodes. First, data of individual electrodes were normalized to the maximum value recorded in the same MEA at the same DIV. Correlation between data of different DIVs in the same electrode (n=236 electrodes per group) was measured by Spearman’s correlation analysis (Fig 6B and Supp Fig 19). Fisher transformation was applied to measure z scores from correlation coefficient values, then z values were used to compare two correlation coefficients between groups [67]. Whether the predefined network structure in 2D and 3D patterned cultures affect the network activity was analyzed by comparing the activity levels in the electrodes that recorded from microwells with 3-edges compared to the electrodes recording from microwell with 4-edges in the center (Fig 6D). We compared the average activity in 3-edge vs. 4-edge nodes in each DIV separately (Supp Fig 13-16) and pooled data from 15 DIV to 45 DIV (Fig 6E-G), based on a paired t-test. A similar analysis was performed for the side and center electrodes of the random networks (Fig 6D).

## Results

### Modular assembly of neuronal networks in 2D and 3D

Morphology of networks during growth and network development in all groups were studied by bright- field microscopy images on different days (Fig 2 and Supp Fig 1-4). In the patterned 2D and 3D cultures, axonal branches extended from microwells into the microchannels between 3 DIV to 5 DIV. At 9 DIV axons in each microwell elongated enough to reach the neighboring modules or extended further and reached to the microwells far away from their originating module (Fig 2F and Supp Fig 1-4). Axonal growth did not simply take a straight path of horizontal or vertical microchannels. Instead, in the crossing points some branches bent and grew perpendicular to their proximal sections (Fig 2F). Neuronal cell bodies remained in the microwell areas, except for some cases in which cells were squeezed into the microchannels in the process of seeding (Supp Fig 2-4). On coverslip cultures, after removing the PDMS devices at 7 DIV network structure gradually distorted by 17 DIV (Fig 2E), and areas between modules were filed with neuronal cell bodies and branches. In contrast, the random networks showed extreme morphological changes in network structure, the position of the cell bodies, and branches between 10DIV and 45 DIV (Supp Fig 1). In the 3D group, bright-field images from z-planes close to the device’s top surface showed the presence of cell bodies and connecting branches (Fig 2G). Nevertheless, the fine details of the 3D network structure were not accessible for bright-field imaging (Fig 2G and Supp Fig 4).

Immunofluorescence images were prepared from networks on coverslips or MEAs. Neurons and astrocytes inside the microwells of 2D and 3D patterned networks were identified (Fig 3 and Supp Fig 6-7). In 2DL cultures, it was possible to remove the PDMS device without causing significant damage to the network structure (Supp Fig 6, Video 2 and Video 3), however in 2DH and 3D engineered networks pealing the PDMS devices affected the circuit structure (Supp Fig 7). Therefore, immunostaining the networks was mainly performed without removing the device (Fig 3A and Supp Fig 7). Nevertheless, without removing the PDMS device, axonal branches inside the microchannels were difficult to be labeled by immunostaining markers (Video 6 and Video 7). Immunostaining of network structure in microwells was straightforward. DAPI-stained nuclei were used to count the average number of cells per microwell (N = 3 cultures and n = 20 microwells per group). The average number of cells per microwell in 2DL, 2DH, and 3D cultures were 16.1 ± 2.31, 37.20 ± 2.98, and 143.91 ± 3.55, respectively. Considering 64 microwells per device, each 2DL, 2DH, and 3D culture on average was composed of 1000, 2380, and 9210 cells, from which at least half of the cells were neurons. The control group (Ctrl) was seeded with an initial density of 100 thousand cells per MEA. These data were later used to measure the percentage of available neurons for recording in each group.

Three-dimensional fluorescent immunohistochemistry and z-stack analysis with confocal microscopy were performed to determine the 3D architecture of the engineered neural networks inside microwells. The overall thickness of the microwells (100 µm) was sliced into 33 z-stack planes with 3 µm distances. Each plane included three channels: DAPI, GFAP, and β-tubulin III. Random and 2D patterned cultures both on the coverslips and MEAs had a maximum thickness of 20 µm while the thickness of the 3D assembled networks was between 70 µm to 100 µm, which was also dependent on the number of cells per well (Fig 3C, Supp Fig 5-7 and Videos 2 to 7). 3D assembled networks in the microwells were connected through axonal branches growing inside the microchannels (Fig 3B, and Video 4 and Video 5). Immunofluorescence images of different z planes confirmed the presence of neuronal cell bodies, axonal branches as well as astrocytes through the whole depth of the microwells (Video 7). The thickness of the axonal tissue inside microchannels was around 5 µm, the same height as the microchannel (Fig 3B-C and Video 4 and Video 5).

### Activity profile of 2D and 3D patterned networks

Activity in all groups appeared as sparse APs at 10 DIV (Fig 4A and Supp Fig 8-11). From 15 DIV we observed first burst activity that reached their peak frequency around 25 DIV (Fig 4A and Supp Fig 8-11). Network activity was evaluated by extracting the AP and burst frequencies recorded from each electrode and averaging it across electrodes of MEAs in the same group at each DIV. The activity of 2DL, 2DH, and 3D patterned networks was compared with activity in random networks of the Ctrl group. From 10 DIV to 25 DIV, we observed an increasing trend of AP frequencies in random and 2D or 3D patterned networks (Fig 4B). Only at 25 DIV, 2DL showed slightly lower AP frequency compared to the random networks (1.93 ± 0.21 AP/s vs. 4.06 ± 0.21 AP/s, p<0.05; Fig 4B). AP frequency in 2DH after 35 DIV was slightly higher than random networks (4.31 ± 0.40 AP/s vs. 2.8 ± 0.20 AP/s, p<0.05; Fig 4B). Starting from 25 DIV, 3D patterned networks showed higher AP frequencies compared to random networks (10.43 ± 0.46 AP/s, p<0.001; Fig 4B).

Burst frequency in the 2DL group was lower than Ctrl group only at 25 DIV (0.04 ± 0.01 bursts/min vs. 0.13 ± 0.02 bursts/min, p<0.01; Fig 4B). 2DH group showed higher burst frequency than Ctrl group at 35 and 45 DIV (0.16 ± 0.02 and 0.15 ± 0.01 bursts/min vs. 0.07 ± 0.01 and 0.11 ± 0.01 bursts/min, respectively, p<0.05; Fig 4B). Burst frequency in the 3D patterned network reached its maximum at 25 DIV (0.35 ± 0.02 bursts/min, p<0.001 vs. Ctrl), followed by a decline at 35 DIV, but was still higher than Ctrl group at 45 DIV (0.17 ± 0.01 bursts/min, p<0.05; Fig 4B). The percentage of APs in the burst represents the tendency of the network for synchronized activity rather than firing individual APs. In 3D networks the percentage of APs in bursts was significantly higher than in other groups from 25 DIV onwards (41.97 ± 1.92 % vs. 23.47 ± 1.83 % in Ctrl group, p<0.001; Fig 4B). 2DL networks showed a lower percentage of APs in a burst at 25 and 35 DIV (2.22 ± 0.46 % and 8.47 ± 1.05 %, p < 0.05). 2DH showed a higher percentage of APs in a burst at 35 DIV compared to the Ctrl group (23.02 ± 2.31 %, p<0.01; Fig 4B). Burst duration in 3D networks was higher than in other groups starting from 15 DIV (0.052 ± 0.01 s vs. 0.039 ± 0.01 s in Ctrl group, p<0.001; Fig 4B). The percentage of APs in bursts and burst duration remained higher after 25 DIV in 3D networks. 2DL networks showed lower burst duration compared to the Ctrl network at 15 and 25 DIVs (p<0.01 and p<0.05, respectively: Fig 4B). Burst duration in the 2DH group showed an increasing trend with culture age (0.042 ± 0.01 vs. 0.03 ± 0.01 in Ctrl group, p<0.05 at 35 DIV).

Each electrode of a MEA can capture the activity of several neurons that are close enough to them. In random networks, most of the neurons are placed in the spaces between electrodes and are therefore not accessible for recording (Fig 2A and Supp Fig 1). Here we tested whether confining the neurons close to the electrodes can improve the readout efficiency. Neural units were extracted by sorting the raw signals in each electrode. The number of detected neural units in each electrode was averaged across different days (15 to 45 DIV) and compared between groups. The average number of detected neural units per electrode in the random networks of the Ctrl group was 3.02 ± 0.10. In 2DL and 2DH groups similar numbers of units per electrode were detected (3.01 ± 0.15 and 3.34 ± 0.14, p=0.99 and p=0.62, respectively; Fig 4D).

In 3D networks each electrode detected average number of 5.49 ± 0.14 neural units (p<0.001 compared to other groups; Fig 4D). Average AP frequency in each neural unit was compared between groups using pooled data between 15 DIV to 45 DIV. These data showed that AP frequency of neural units in 2DL networks was slightly lower than Ctrl and 2DH groups (0.97 ± 0.08 vs. 1.13 ± 0.05 and 1.13 ± 0.09 AP/s, p < 0.05; Fig 4D). In the 3D network, a higher AP frequency was recorded compared to the random or 2D patterned networks (1.81 ± 0.08, p<0.0001; Fig 4D). The percentage of available neurons for recording in each culture was measured based on the total number of neurons per culture and the total number of detected units per electrode. In the Ctrl group, only 0.21 ± 0.01 % of neuronal activity was detectable which means on average 210 neurons out of 100,000 seeded cells were functionally detected by 59 electrodes of an MEA. The highest percentage of available neurons for recording was acquired by seeding them in the lowest density in the 2DL group (18.71 ± 0.95 %, p<0.001 *vs.* other groups; Fig 4D). 2DH and 3D cultures showed a higher percentage of available neural units for recording compared to the Ctrl group (8.97 ± 0.38% and 3.84 ± 0.10%, respectively, p < 0.001; Fig 4D). The percentage of available neurons for recording in the 3D group was lower than the 2DH group (p < 0.001; Fig 4D).

### Functional connectivity maps in 2D and 3D modular networks

Functional connectivity between two modules in patterned networks was extracted based on cross-correlation analysis between pairs of corresponding electrodes recording from those two modules. Connectivity maps were plotted based on the presence of a functional connection between modules if the value of the correlation coefficient was higher than 0.25. The whole network functional connectivity map was plotted based on detected connectivity between each pair of electrodes. The total number of detected connections per electrode was depicted by the size of the electrode and the strength or weight of connection was illustrated by the thickness of the connecting lines (Fig 5A-B). Representative whole network connectivity maps of a selected MEA culture from each group at 15 DIV to 45 DIV are shown in Fig 5A. Examples of detailed changes and dynamic complexity of the connectivity maps by culture age have been displayed in Video 8.

**Fig 5:**
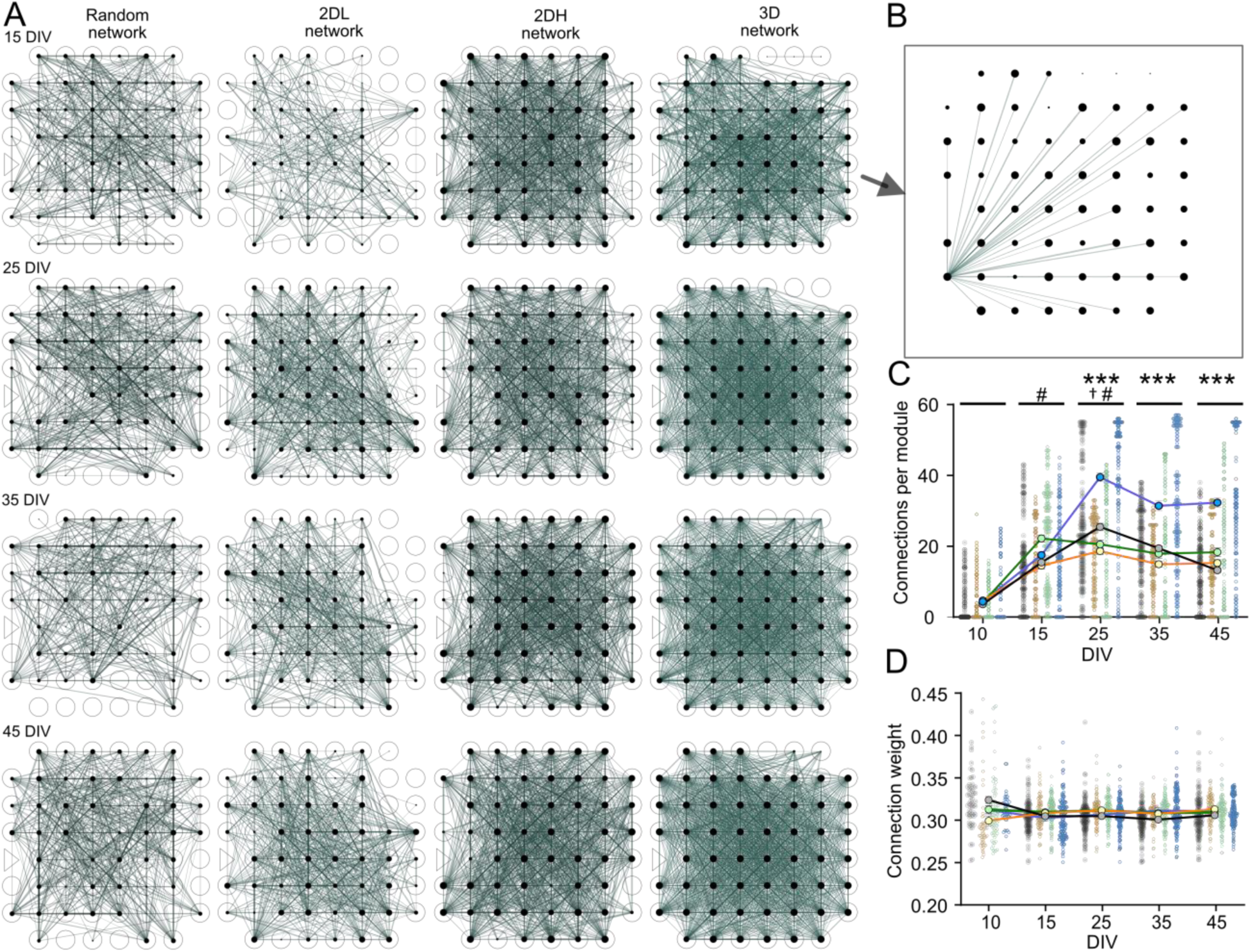
Functional connectivity maps in random and modular networks. **A**) Functional connectivity map in a selected MEA from each group at different DIVs. In each plot, the size of the electrode (black circle) represents the number of connections per module. Line thickness represents the weight of the connection. **B**) Representative image of an electrode with all connections at 15 DIV from 3D network. The size of electrodes represents number of connections. **C** and **D**) Number and strength of connections at different DIVs. Each point represents the average number of connections or connection strength per electrode (N=4 MEA per group and n=236 electrodes per group). Data has been presented as Mean ± S.E.M. Data between groups were compared using mixed-effects analysis followed by Tukey’s multiple comparison test. 2DL *vs.* Ctrl († p < 0.05), 2DH *vs.* Ctrl (# p < 0.05,), and 3D *vs.* Ctrl (*** p < 0.001).

The number of connections per module, or electrode, increases by culture age between 10 DIV and 25 DIV for random and 2D or 3D patterned circuits and remained relatively stable afterward (Fig 5). At 25 DIV, the average number of connections per module in 3D modular networks was higher than random, 2DL and 2DH networks (39.51 ± 1.27 connections, vs. 25.47 ± 1.00, 18.63 ± 0.90 and 20.55 ± 1.24, respectively, p < 0.001, Fig 5C). The number of connections per module in 3D networks remained higher than other groups for the rest of the study (p<0.001, Fig 5C). We did not observe a significant difference in the number of connections per module between 2D patterned networks and random networks (Fig 5C). Despite the highly dynamic nature of individual connection weights, no significant difference was observed comparing the average connection strength along with culture age and as well between groups (Fig 5D).

### Impact of structure on function in 2D and 3D patterned network

Besides the differences in the total number of neurons, two main structural differences between patterned and random networks were (1) limited changes in the position of the neurons and neurites by culture age, and (2) modular network structure with 64 nodes, each with 3 or 4 edges, in patterned cultures (Fig 1 and Fig 6).

**Fig. 6.**
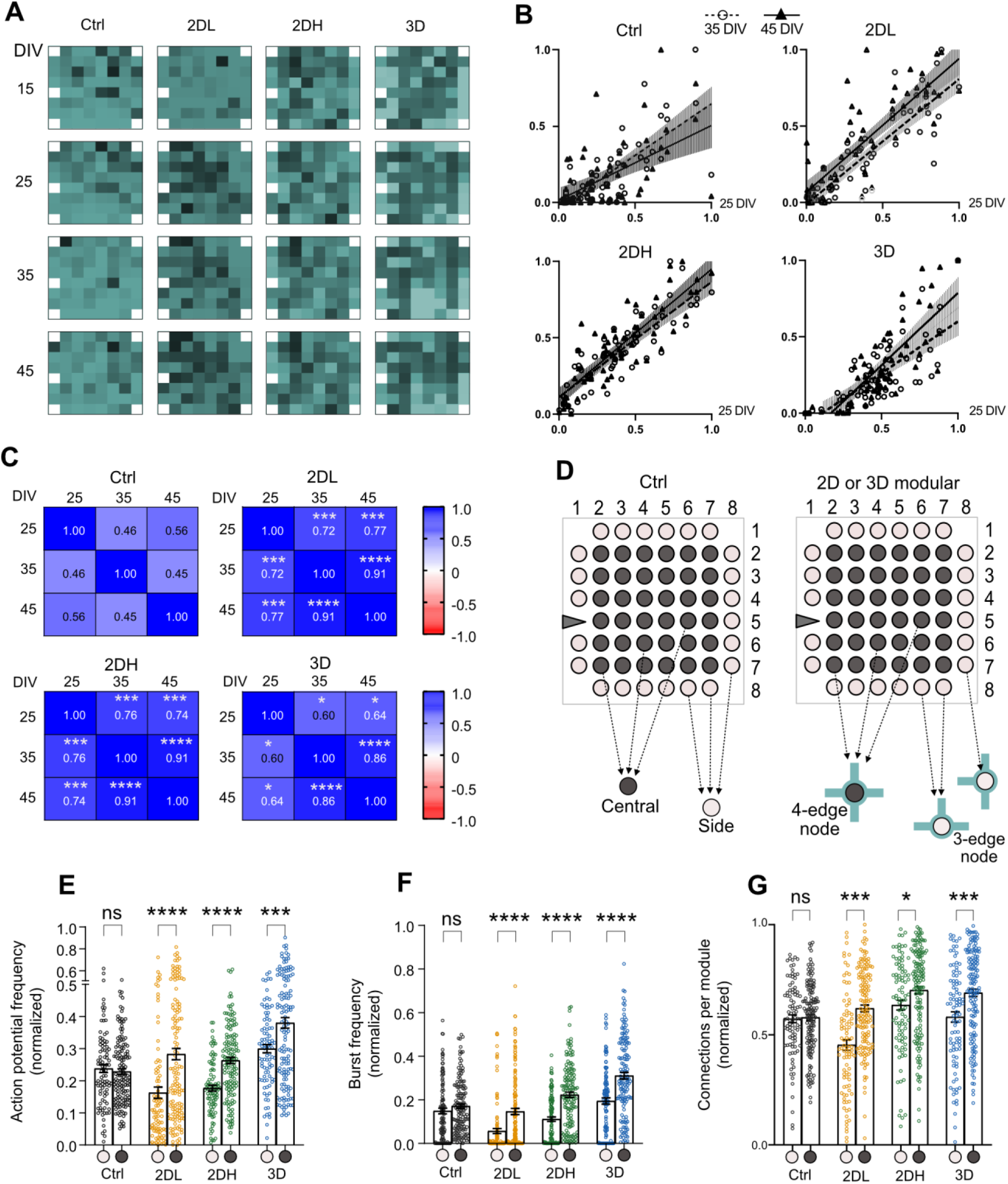
Impact of network structure on activity. **A**) Heatmap of normalized activity in a selected MEA from each group in different DIVs. In each DIV, AP frequency in each electrode of MEA was normalized to the maximum AP frequency recorded in that MEA. Normalized activity in each electrode is tracked over time. **B**) Correlation of AP frequency data between 25 vs. 35 DIVs (circles and dashed line) and between 35 vs. 45 DIVs (triangle, straight line) in a selected MEA from each group. C) Heatmap table with correlation coefficient (CC) values of Spearman’s r-value between DIVs of the same group (N=4 MEA and n=236 electrodes per group). Fisher r to z transformation followed by student t-test to compare CCs between patterned networks with the random network (* p < 0.05, ** p < 0.01, *** p < 0.001 and **** p < 0.0001). **D**) Each electrode, based on its position, records from side modules, or nodes, with 3-edge (n = 23 per MEA) or from central nodes with 4-edge (n = 36 per MEA). **E-G**) Normalized AP and burst frequency and number of connections per module in 3-edge vs. 4-edge nodes of the same group. Data pooled from 15 DIV to 45 DIV recordings. In each node, data was normalized to the maximum value of the same MEA and then averaged across DIVs. Each point represents pooled data from one module or electrode. Data compared between 3-edge vs. 4- edge nodes of the same group using paired t-test (*** p < 0.001, and **** p < 0.0001). Comparisons between 3-edge *vs.* 4-edge nodes at specific DIVs have been summarized in the Supp Fig 13-18 including other burst features and connectivity strength.

### Spatial distribution of activity and functional connectivity parameters over time

To understand whether a stable network structure between 15 DIV to 45 DIV affected the activity profile we tracked the activity features and functional connectivity parameters in each microwell, node, or electrode over time in patterned cultures *vs.* random networks. To make it comparable between DIVs and MEAs, in each DIV values of individual electrodes were normalized to the maximum value obtained in the same MEA culture on that day. Then we determined if normalized AP frequency, burst features, and number and strength of connections per module in individual electrodes, or microwells, change at different DIVs. If an individual electrode or node shares the same ratio of overall network activity or functional connectivity in two different DIVs there should be a positive correlation between days, indicating that activity or connectivity pattern is stable over time in different parts of the network. Meaning that, despite changes in overall network activity between two DIVs, each node shares a comparable percentage of activity in both DIVs, and network activity is spatially stable. Conversely, if an electrode does not share a comparable ratio of overall network activity at two different DIVs, no correlation between DIVs should be observed, which indicates that network activity shifts considerably over electrodes and is not spatially consistent between DIVs. These data have been summarized in Fig 6A-C and Supp Fig 19.

An example spatial distribution of AP frequency over time is shown in Fig 6A which displays normalized AP frequency in electrodes of a selected MEA in each group at 15 DIV to 45 DIV. Compared to random networks which showed a distinct spatial distribution of activity at different DIVs, patterned 2DL, 2DH, and 3D networks showed consistent spatial distribution of activity at different DIVs, except at 15 DIV (Fig 6A). For instance, in a 2DL culture electrode 44, or microwell 44, is among the most active electrodes at 25 DIV, the same electrode remained among most active electrodes on 35 DIV and 45 DIV (Fig 6A). This was statistically confirmed by the positive correlation between normalized values of AP frequency at different DIVs in pooled data from MEAs of the same group (N = 4 MEAs and n = 236 electrodes per group; Fig 6B- C). Values of the correlation coefficient (Spearman’s r) between 25 DIV, 35 DIV, and 45 DIV of the same group are summarized in Fig 6C. Correlation coefficients (CCs) between 25 DIV and 45 DIV were 0.56 for Ctrl, 0.77 for 2DL, 0.74 for 2DH and 0.64 for 3D groups (Fig 6C), and between 35 DIV and 45 DIV we observed r-values of 0.45, 0.91, 0.91 and 0.86 for Ctrl, 2DL, 2DH, and 3D groups, respectively (Fig 6C). Fisher r to z transformation was applied to compare the CCs between groups at different DIVs (Fig 6C). CC of AP frequency data between 25 DIV, 35 DIV, and 45 DIV in 2DL and 2DH groups was significantly higher than random networks (p<0.001, Fig 6C). 3D modular networks showed higher CC between 25-35 DIV, 25-45 DIV and 35-45 DIV (p<0.05, p<0.05 and p<0.0001 vs. Ctrl group, Fig 6C).

The same analysis was performed for normalized burst frequency, burst duration, and percentage of APs in the burst. These showed comparable results (Supp Fig 19). In 2D modular networks, all burst features showed a positive correlation between 25, 35, and 45 DIVs, though CC between 35 and 45 DIVs was greater than 25 and 45 DIV (Supp Fig 19). In 3D modular networks burst frequency and percentage of APs in the burst showed a positive correlation between 25 and 45 DIV (r>0.64) and 35 and 45 DIVs (r>0.79). Burst duration in the 3D patterned group was only correlated between 35 and 45 DIVs (r=0.84; Supp Fig 19). All burst features of random networks also showed a positive correlation with lower CC values between 25 DIV to 45 DIV (r<0.60 for all comparisons, Supp Fig 19). For all burst features CCs in 2DL and 2DH groups were significantly greater than random networks (p-values have been summarized in Supp Fig 19). Burst frequency and percentage of APs in a burst in 3D patterned networks showed stronger CC compared to random networks (between 25 DIV to 45DIV, Supp Fig 19). Correlation studies of burst duration between 35 DIV and 45 DIV in the 3D group showed higher CCs compared to the random networks (p<0.0001), while CC between 25 DIV and 45 DIV was lower than random networks (p<0.0001).

We also probed the spatial distribution of functional connectivity parameters over time. CC values for number of connections per module at 25 vs. 45 DIV and 35 vs. 45 DIV in 2DL patterned networks (r=0.52 and 0.74), in 2DH networks (r= 0.64 and 0.76), and in 3D networks (r=0.67 and 0.71), were significantly greater than random networks (r=0.24 and 0.49, p-values have been summarized in Supp Fig 19). Strength of connection was not correlated between 25 vs. 45 DIVs and 35 vs. 45 DIVs in random networks and 3D patterned circuits (r=0.42 and 0.45, and r=0.05 and 0.55, respectively; Supp Fig 19). In 2DL, 2DH and 3D modular networks CCs between 35 vs. 45 DIV were higher than random networks (r=0.69, 0.67, and 0.55 vs. r=0.45 in Ctrl group).

### Effect of physical edge numbers on activity and functional connectivity profile in modular networks

Whether predefined network structure affects the activity and functional connectivity profile in different modules was tested by probing the activity and connectivity parameters in microwells that connected through 3 vs. 4 microchannels or physical edges. Based on design configurations, activity in 3-edge microwells was recorded by electrodes at the border of the sensing area, and activity of 4-edge microwells was monitored by electrodes in the center (Fig 6D). We also compared the same parameters in the random networks between electrodes located in the side vs. center of the sensing area (Fig 6B). In the random network, we did not observe a significant difference in AP or burst frequency of electrodes that are in the edge of the configuration compared to electrodes located in the center (p=0.92 and p=0.56, respectively, Fig 6E-F and Supp Fig 13-14). In 2DL, 2DH and 3D patterned networks AP frequency of 4-edge nodes were significantly higher than 3-edge nodes (p<0.0001, p<0.0001 and p<0.001, respectively; Fig 6E and Supp Fig 13). In 2DL, 2DH, and 3D cultures burst frequency in 4-edge nodes was also higher than 3-edge nodes (p<0.0001; Fig 6F and Supp Fig 14). Other burst features including the percentage of APs in the burst and burst duration were also significantly higher in the 4-edge nodes compared to the 3-edge nodes in the 2D and 3D patterned cultures (Supp Fig 15 and Supp Fig 16).

Pooled data from 15 to 45 DIV in the patterned 2DL, 2DH, and 3D networks revealed a greater number of functional connections for 4-edge modules compared to the 3-edge modules (p<0.001, p<0.05, and p<0.001, Fig 6G). The same results were observed if 4-edge and 3-edge modules were compared at individual DIVs (Supp Fig 17). In random networks electrodes located in edge *vs.* the ones in the central regions of the sensing area showed a similar number of connections per electrode (p=0.99; Fig 6G and Supp Fig 18). The strength of connections in 4-edge modules of 2DL, 2DH, and 3D networks was slightly higher than 3-edge modules, however, the difference was not statistically significant (p=0.09, p=0.92, and p=0.07 at 35 DIV; Supp Fig 18).

## Discussion

Neuronal networks with reproducible functional attributes can serve as optimized *in vitro* platforms to model functional phenotypes of neurological disorders, screening their response to treatment candidates, as well as addressing basic questions such as learning and plasticity [68–70]. Functional data of the randomly assembled networks often show large variation between cultures and among different ages of the same network [27]. This is mainly related to the arbitrary structure of the network in each culture and the constant shift in the position of the soma and axon with culture age [21,27]. One of the main advantages of the patterned neuronal circuits, therefore, is their predefined structural composition that remains stable over months [31,41,71,72]. Therefore, we studied the reproducibility of the structure-dependent functional output in 2D and 3D modular cortical networks.

To explore the functional characteristics of modular networks, several labs have engineered 2D neural circuits on planar MEA electrodes [31,39,44,73,74]. Previously studied configurations of patterned circuits differ mainly with respect to the number of edges, or connections, per module from linear structures to triangular, square, and hexagonal structures with 2, 3, 4, and 6 connections per module, respectively [31,39,44,74–76]. Here, we designed and compared 3-edge and 4-edge modules in the context of the same network (Fig 1). Such networks with a heterogeneous number of edges per module are closer to *in vivo* circuits and offer an optimized context to compare the impact of structure on function. The same physical design allowed us to assemble 3D modular networks aligned with MEA electrodes (Fig 1). Self-assembled 100 µm thick 3D structure in each node was composed of neurons and astrocytes that was supported by PDL-coated microwells. Previous models were limited to engineering uniform and randomly connected 3D neural circuits on MEA substrates [53,57,59]. To mimic modular brain circuits we designed, for the first time, a 3D modular network on planar MEA electrodes with long-term access to network electrophysiology data.

Modular engineering improved the electrophysiology readout efficiency by confining the network to the recording sites (Fig 3). In 2D patterned networks, the percentage of available neuronal units for recording was at least 40 times more than with random networks (Fig 4D). To improve the electrophysiology readout efficiency from random networks, high-density MEAs with a higher number of electrodes and smaller electrode pitch are beneficial [15,77]. This technology could be combined with our modular microchannel design. Even though in our 3D modular model the readout efficiency was 10 times higher than in random 2D networks, planar electrodes allow recording only from the bottom side of the network [53]. This challenge could be resolved by recently developed 3D MEA systems that offer multisite recording from 3D neural culture microsystems and organoids [55,59,78,79]. Nevertheless, the feasibility of constructing functional 3D modular networks on currently available models of 3D MEAs needs to be tested.

Previously, some labs tried to compare the activity of 2D modular and random networks derived from the rodent hippocampus, which showed distinct results between the groups [29,74,75,80]. For instance, Marconi et al. recorded a higher rate of AP and burst activity in patterned hippocampal networks [29], while Boehler et al obtained opposite results [80]. Here, in cortical cultures, average activity in each electrode was slightly lower in 2DL patterned networks and slightly higher in 2DH patterned networks (Fig 4B). However, the AP firing rate of individual neuronal units was in the same range for 2D patterned and random networks (Fig 4D). 3D patterned cortical networks, on the other hand, represented elevated activity levels compared to the 2D networks. Strong burst features including longer bursts in our 3D modular networks are comparable to the burst activity recorded from random 3D engineered cortical networks and spheroids [53,58]. Longer bursts are the functional signature of the developing neonatal rat cortex [81].

Burst activity in random cortical networks develops between second and third week and reaches its peak around 21 DIV [30,61,82], which indicates development and maturation of the synapses [83]. In random and modular cortical networks investigated here, functional connectivity data and burst features showed comparable trends. Functional connections increased within the second and third week (10-25 DIV) in random and patterned networks and remained constant for the rest of the study (Fig 5C). Immunostaining results have previously shown that the number of synaptic connections in 2D patterned and random networks increase within the second and third week and remain constant thereafter [29,75]. Accompanied by stronger activity levels a higher number of connections per node was detected in 3D modular networks (Fig 4 and Fig 5). Recently, Tibau et al. used PDMS stencils to culture aggregated and semi-aggregated cortical networks and compared their functional trait with homogenously distributed networks using calcium imaging [60]. Aggregated cortical networks showed a higher number of connections compared to semi-aggregated and homogenously distributed networks [60]. These data, together with our results suggest that spatial proximity of neurons in aggregates or 3D assembled modules substantially enhances the functional connectivity of cortical networks and therefore their synchronized burst activity.

The spatial distribution of neurons in the developing brain is established by well-regulated neuronal motility and migration [84,85]. *In vitro* network structure is also altered by neuronal tendency to move and relocate [27,86], as we observed in our random networks on MEA substrate (Supp Fig 1). The activity of a moving neuron might be temporarily detectable at a specific day, when it is close enough to a recording electrode, while it can disappear a few days later [27]. In a recent study, Habibey *et al.,* showed that along with network morphology changes by culture age neuronal cells position shifted more than 200 µm in average [27]. Therefore, neuronal motility in random networks can affect the reliability of long-term electrophysiology data. This issue is addressed, to some extent, by densely packed electrodes of high-density MEAs [25,27,87]. Nevertheless, in high-density MEAs detectable activity of a moving neuron can shift between electrodes at different days. In modular networks, the neuronal movement was essentially limited to a microwell and the corresponding electrode. This assured each electrode is recording from the same part of the network or same group of neurons at different DIVs. To determine if the stability of the network affects the consistency of long-term electrophysiology data, we tracked the spatial distribution of activity in individual electrodes over weeks. Compared to the random networks, modular 2D and 3D networks established a more consistent spatial distribution of AP frequency, burst features, and connection numbers (Fig 6 and Supp Fig 19). This consistency in 2D modular networks was higher than 3D networks, most likely related to the availability of neuronal movement in vertical axes. Reliable long-term functional data derived from the stable structure of the modular circuits can overcome the hurdles of tracking the functional phenotype of diseased networks *in vitro* [68].

Based on the modern theory of structure-function relationship in the brain, network structure profoundly affects its function [8]. Here the impact of network structure on function was further assessed by probing the functional features in nodes with a different number of physical connections. Boehler et al. have previously shown that overall activity in modular networks increases by including more edges per node [80]. In our model, modules with 3 or 4 edges in the same network represented different functional phenotypes: elevated activity levels, longer burst duration, and higher number of functional connections in the 4-edge nodes vs. 3-edge nodes. Nodes with 4-edges were physically central compared to the 3-edge nodes on the side of the network structure. In the brain, microcircuits with nodes located in the center of the network structure tend to be also functionally central nodes with a higher number of connections [8,88]. The central electrodes of the random networks, however, did not show a higher activity or connectivity compared to the side electrodes. Geometry and number of physical connections in 4-edge nodes resulted in elevated activity and connection number. Even though, the theory of structure-function relationship is still in its infancy [8], engineering functionally predictable neuronal circuits *in vitro* offers a tool with advanced potentials for the neuroscience community.

## Conclusion

By engineering 2D and 3D modular networks of cortical neurons on MEAs, we aimed to mimic the of modularity in brain networks [3]. Each 2D or 3D assembled module is composed of astrocytes and neurons connected through microchannels to the adjacent modules. Assembling the 2D and 3D networks in PDMS microwells confined neurons to the recording sites that improved the readout efficiency and preserved network structure over weeks. Patterned 2D circuits were created by a considerably lower number of neurons, however, represented comparable activity profile and functional development compared to 2D random networks. Modular 3D networks with profound activity features such as prolonged bursts, and higher density of connections recapitulated key functional features of developing neonatal cortex. Modules with a different number of physical connections revealed the impact of engineered network structure on functional output. Stable network structure in modular cultures established a consistent spatially distributed activity which offers great potential to develop more reliable functional readout tools.

PDMS microstructures can be designed to generate and test more complex neuronal circuits in 2D or 3D. This approach can be integrated with technological advances of high-density MEAs and 3DMEAs to extend the competence of the patterned circuits for deciphering the functional features of developing networks. This approach will serve both basic biomedical research as well as studying pathophysiological phenotypes of developing networks derived from human patients. We believe that progress in engineering neural circuits parallel to self-assembled organoid systems can provide a crucial step to promote the *in vitro* systems as a complementary platform to *in vivo* research within the neuroscience community.

## Author contribution

Conceptualization: RH; Methodology: RH, SL; Software: RH, JS, RL, FS, SL; Validation: RH, JS; Formal analysis: RH, RL, SL, JS; Investigation: RH, SL; Resources: RH, SL; Data curation: RH; Writing-Original draft: RH; Writing- Review & Editing; RH, JS, SL; Visualization: RH, SL; Supervision; RH; Project administration: RH, SL; Funding acquisition: RH, SL.

## Acknowledgments

RH acknowledges funding from Volkswagen Foundation (Freigeist–A110720). JS acknowledges support by the Joachim Herz Foundation.

## Notes

### Competing Interest Statement

The authors have declared no competing interest.

