## Supplementary Figures and Text for "Microengineered 2D and 3D modular neuronal networks represent structure-function relationship"

**Title:**

### Video Files:

Video 1: Animated steps of integrating microfluidic and MEA devices and cell seeding, consisting of aligning the PDMS microdevice on the electrodes of the MEA, placing the PDMS mask, loading cell suspension on the device, mask removal, filling the MEA chamber with media, and sealing the chamber with a custom-made PDMS lid.

Video 2: Three-dimensional view of 2D random networks and patterned 2DL and 2DH cultures on MEA or coverslip. All videos were prepared from at least 33 confocal z-series of merged channels (DAPI, β-tubulin III, and GFAP). Z-axis values represent the culture thickness in µm. Microscopy images from 2DH network on MEA were prepared using an inverted microscope that shows position of the electrodes.

Video 3: Animation showing confocal *z*-series imaging of four adjacent modules in 2DL patterned network form MEA electrodes upwards.

Video 4: Three-dimensional rotation of reconstructed images from 9 adjacent modules of a 3D patterned network on a coverslip. In total 33 confocal z-series of merged channels (DAPI, β-tubulin III, and GFAP) were used to create the 3D view of the network. Z-axis value represents the culture thickness in µm.

Video 5: Animation showing confocal *z*-series imaging of 9 adjacent modules in 3D patterned network of Video 4. All channels and the merged image are represented separately.

Video 6: Three-dimensional rotation of reconstructed images from 9 adjacent modules of a 3D patterned network on the MEA substrate. In total 33 confocal z-series of all channels separately (DAPI, β-tubulin III, and GFAP) and their merged image were used to create the 3D view of the network. Z-axis value represents the culture thickness in µm.

Video 7: Animation showing confocal *z*-series imaging of 9 adjacent modules in 3D patterned network of Video 6 on MEA substrate. All channels and their merged images are represented separately. A magnified view of a 3D network structure inside a microwell is shown in the upper right corner.

Video 8: Changes in functional connectivity maps over time in a selected MEA from each group. The size of each black circle represents the average number of connections per electrode and line thickness represents the weight of the connection.

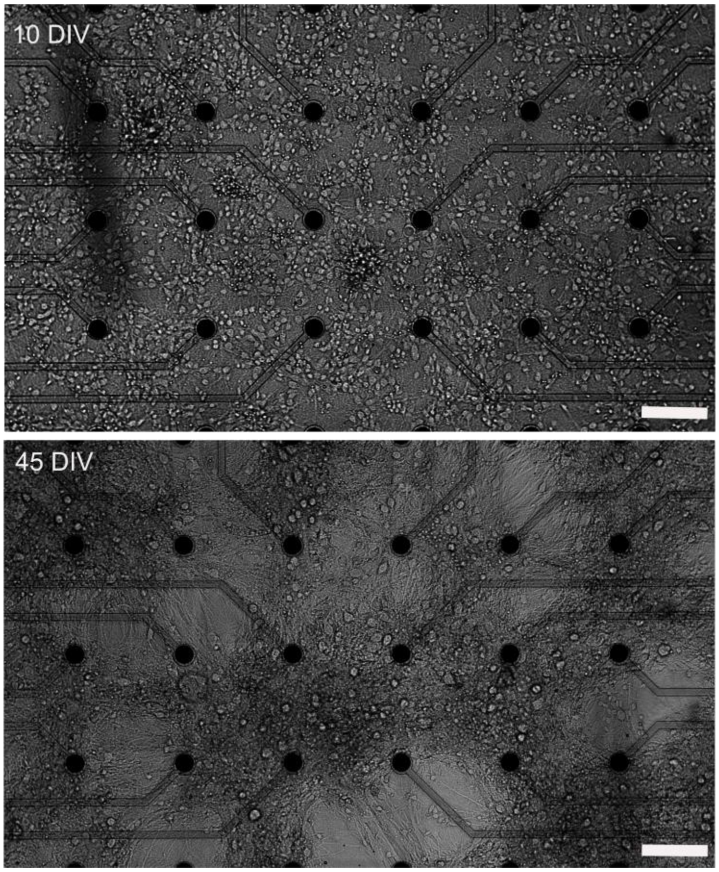


**Supp Fig. 1 Random network morphology in the control group (Ctrl) at 10 DIV and 45 DIV.** The network structure and position of the neurons drastically change by culture age. Scale bar = 100 µm.

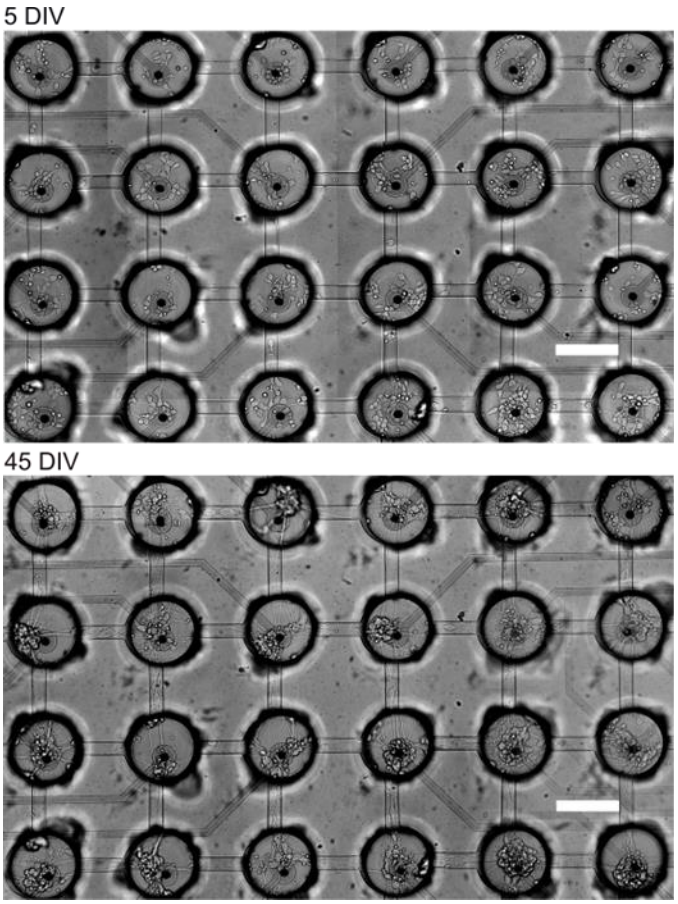


**Supp Fig. 2 Network morphology in low-density 2D patterned (2DL) group at 5 DIV and 45 DIV.** PDMS structures confine the small-world neuronal modules to each microwell while communicating through connecting microchannels to other modules. Scale bar = 100 µm.

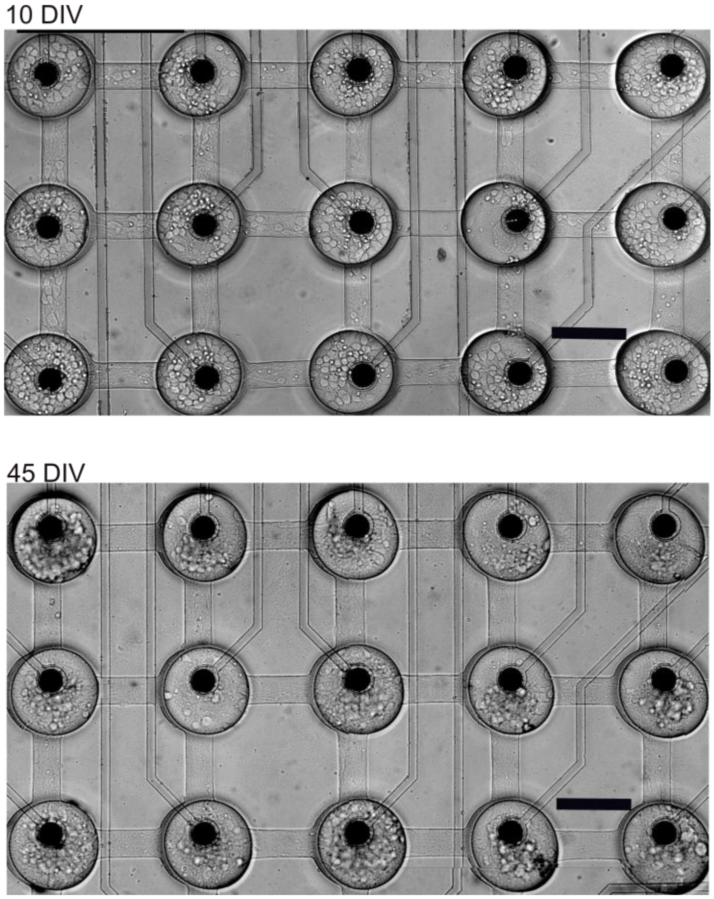


**Supp Fig. 3 Network morphology in high-density 2D patterned (2DH) group at 10 DIV and 45 DIV.** Scale bar = 100 µm.

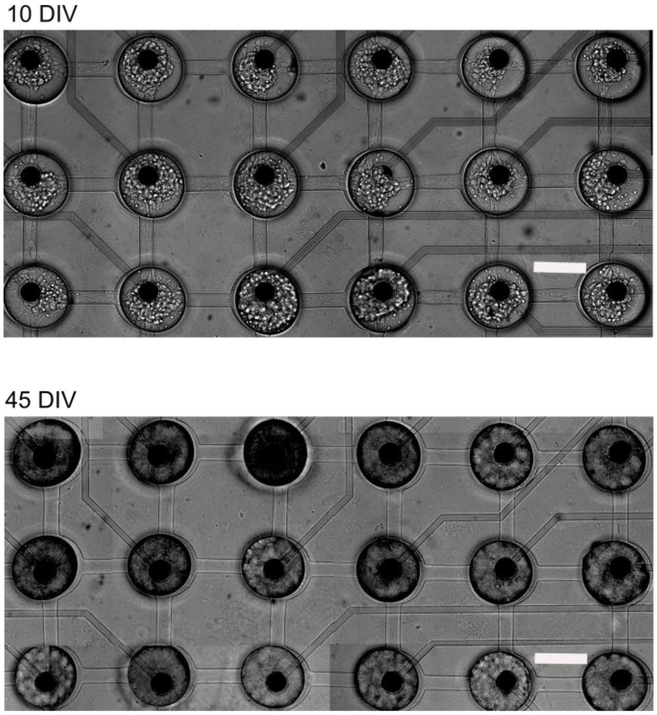


**Supp Fig. 4 Network morphology in 3D patterned group at 10 DIV and 45 DIV.** Scale bar = 100 μm.

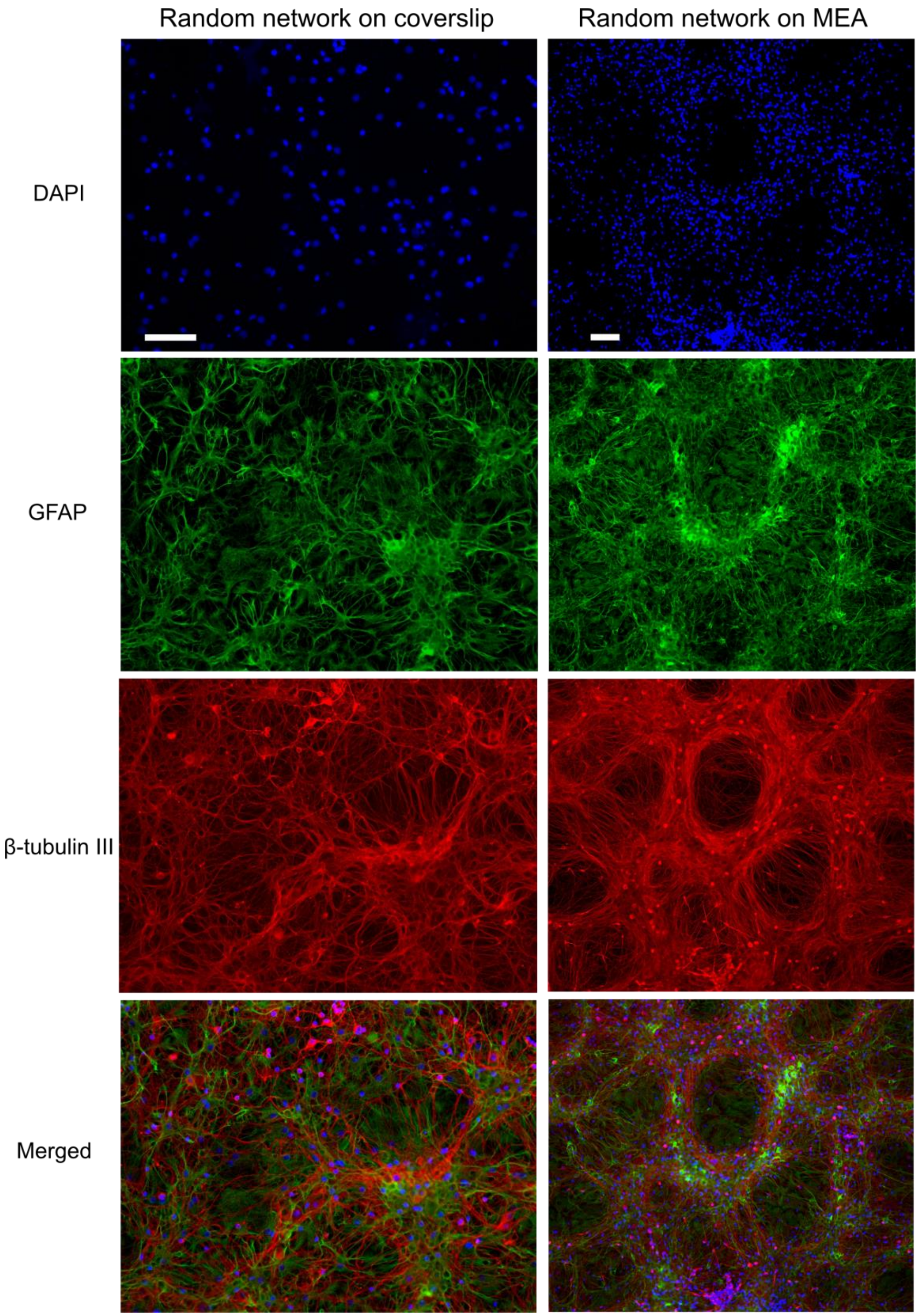


**Supp Fig. 5 Immunofluorescence images of network morphology in control group (Ctrl) at 45 DIV.** images have been prepared from cultures on coverslips (left) and MEA (right) using an inverted confocal microscope (Nikon A1+). Scale bar = 100 µm.

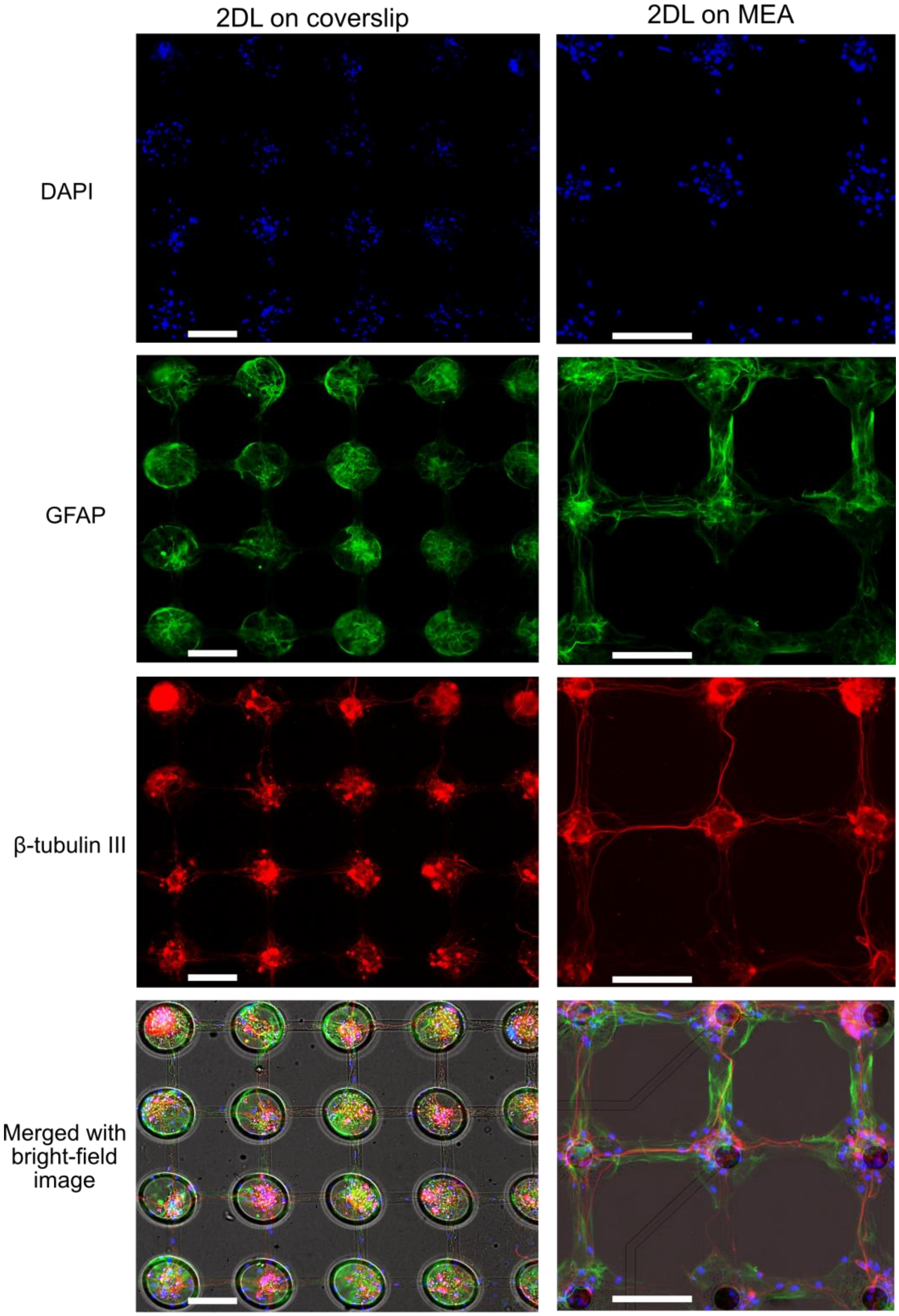


**Supp Fig. 6 Immunofluorescence images of network morphology in low-density 2D patterned culture (2DL) at 45 DIV.** Images have been prepared from cultures on coverslips (left) and MEA (right) using an inverted confocal microscope (Nikon A1+). PDMS device has been removed before immunostaining. Scale bar = 100 μm.

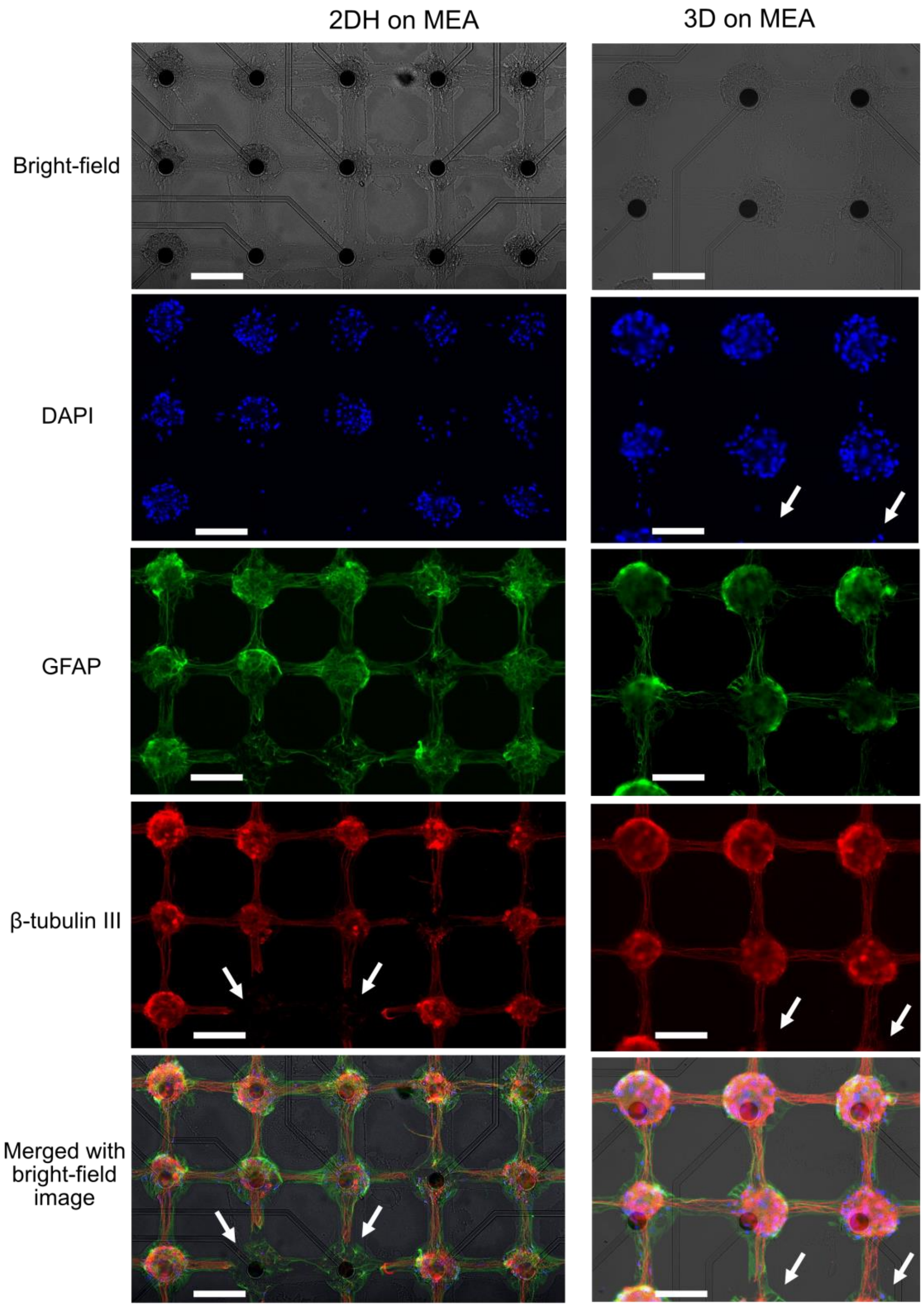


**Supp Fig. 7 Immunofluorescence images of network morphology in high-density 2D patterned (2DH) and 3D patterned culture at 45 DIV.** Images have been prepared from cultures on MEA using an inverted confocal microscope (Nikon A1+). PDMS device has been removed before immunostaining that affected some parts of the network indicated by white arrows. Scale bar = 100 µm.

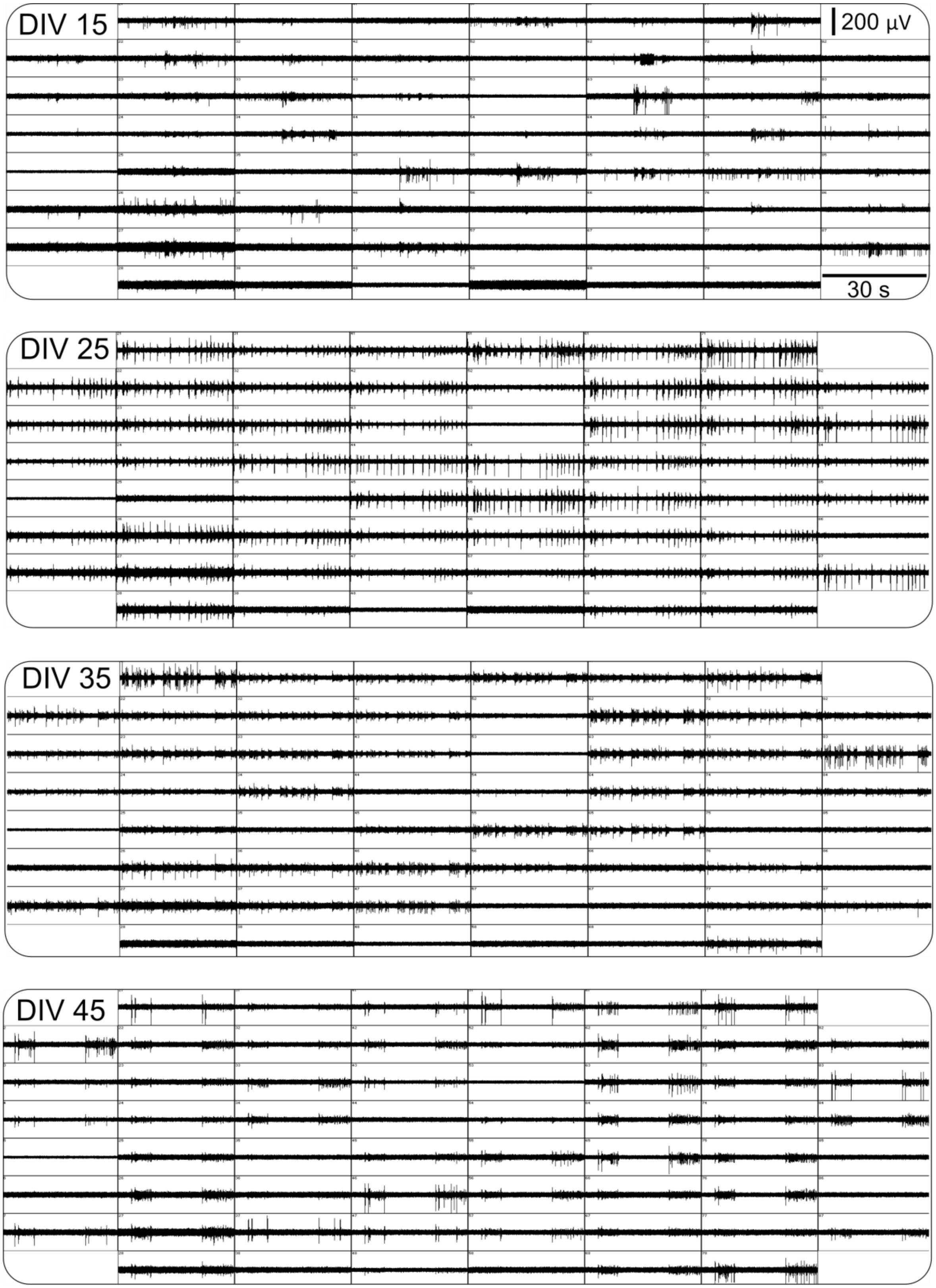


**Supp Fig. 8 Activity profile of a selected random culture in Ctrl group at different DIVs. Each window represents 30 s activity profile at recorded by one electrode.**

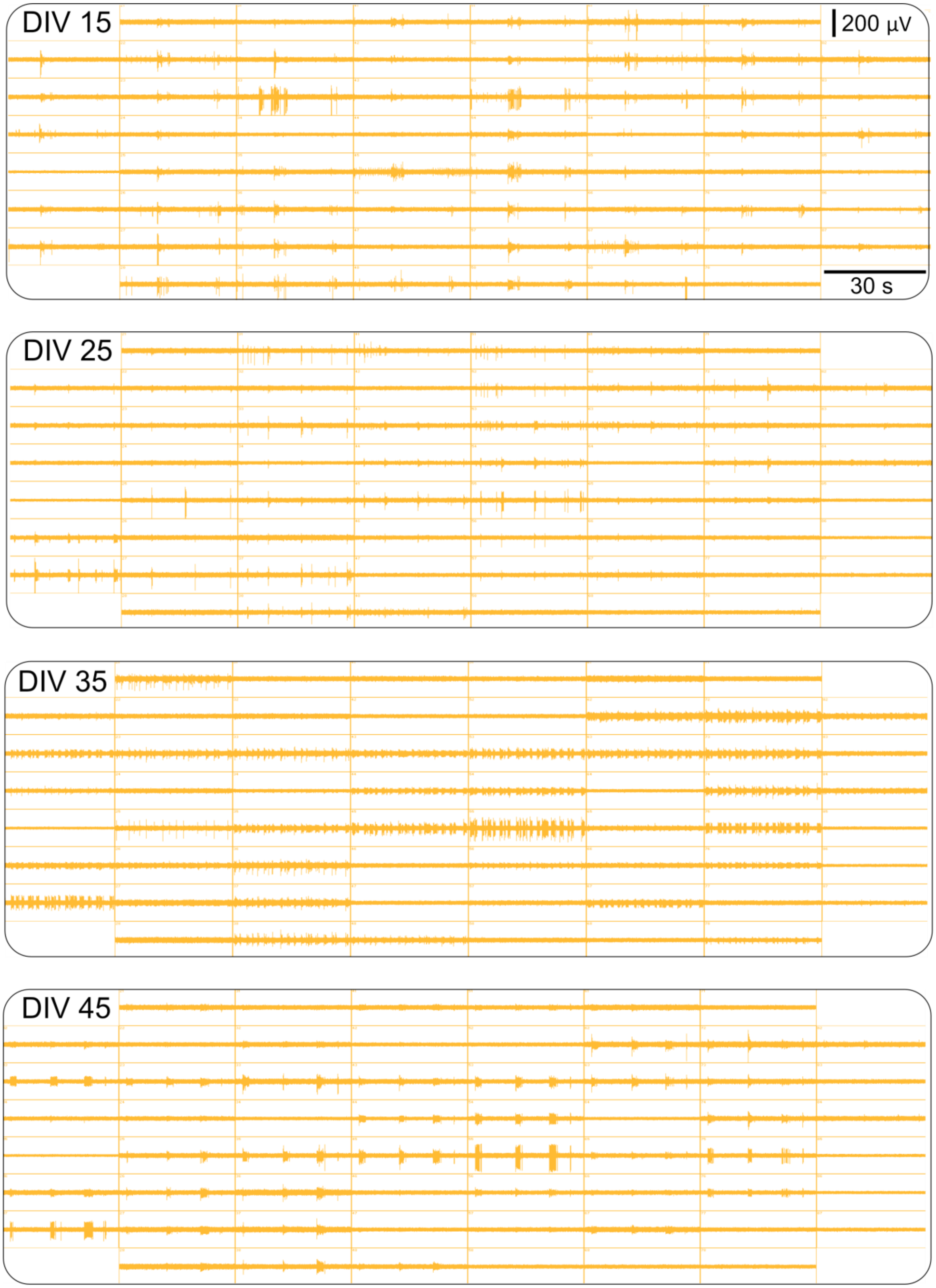


**Supp Fig. 9 Activity profile of a selected low-density 2D patterned network (2DL) at different DIVs. Each window represents 30 s activity profile at recorded by one electrode.**

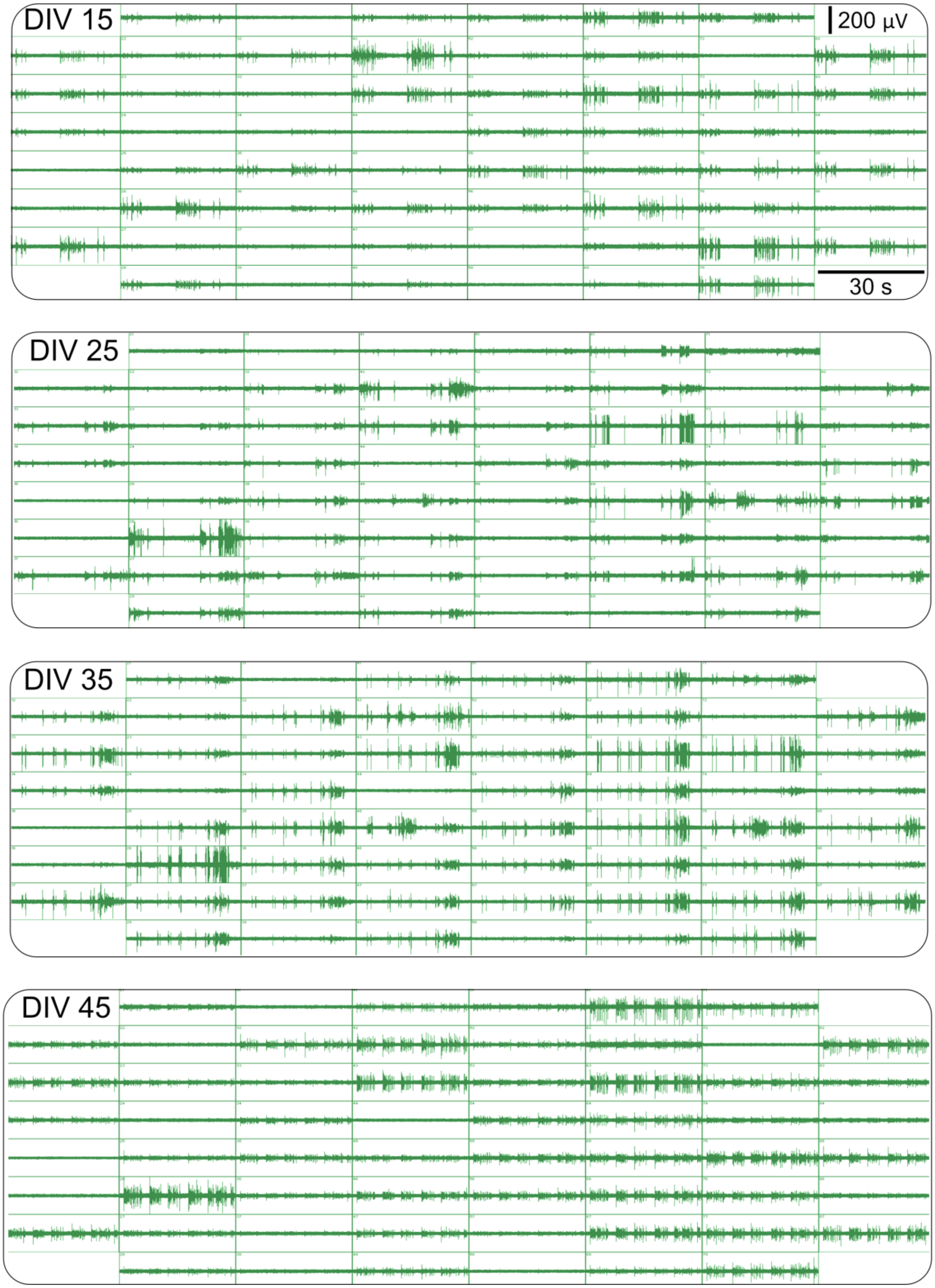


**Supp Fig. 10** **Activity profile of a selected high-density 2D patterned network (2DH) at different DIVs. Each window represents 30 s activity profile at recorded by one electrode.**

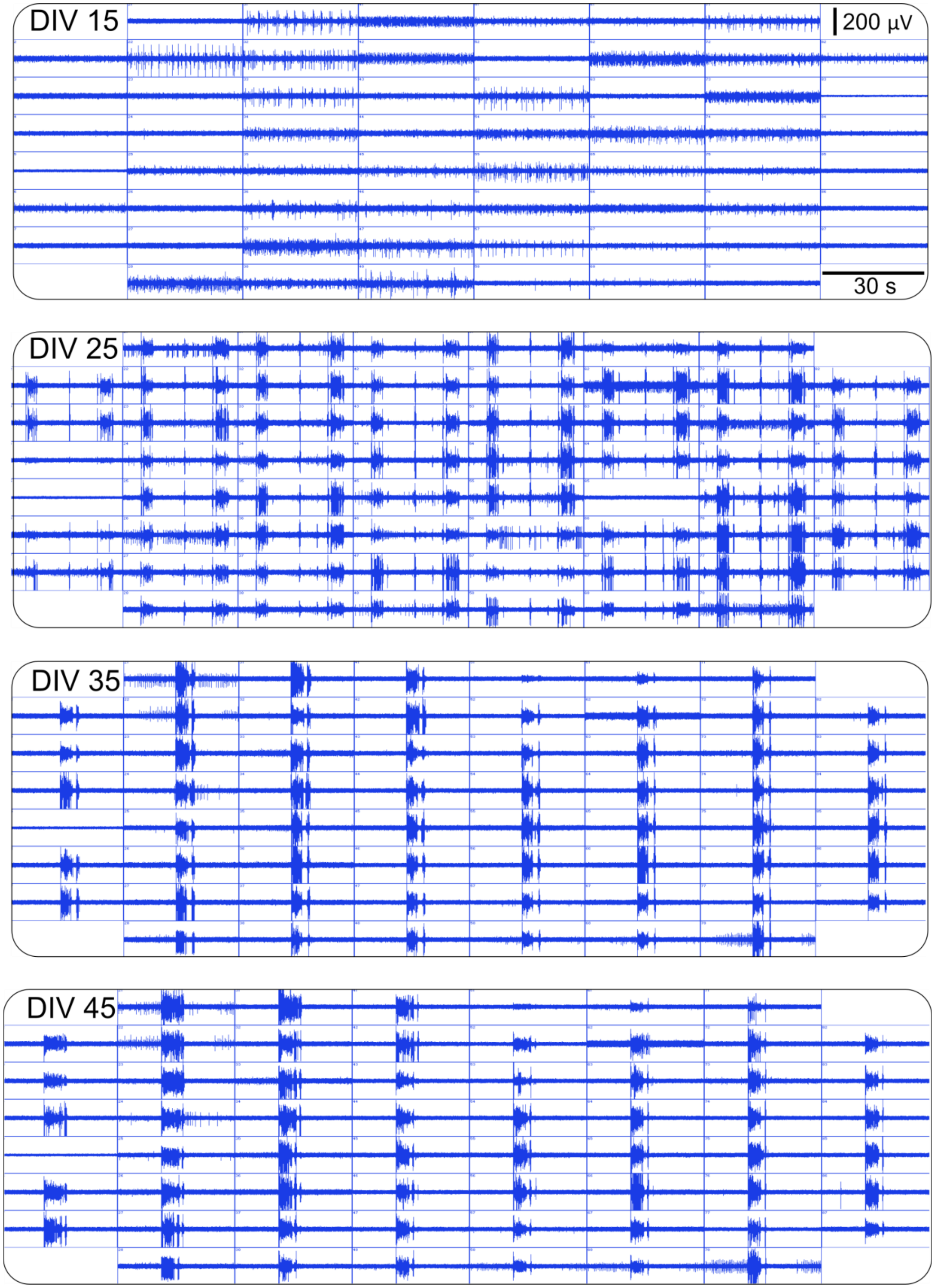


**Supp Fig. 11 Activity profile of a selected 3D patterned network at different DIVs. Each window represents 30 s activity profile at recorded by one electrode.**

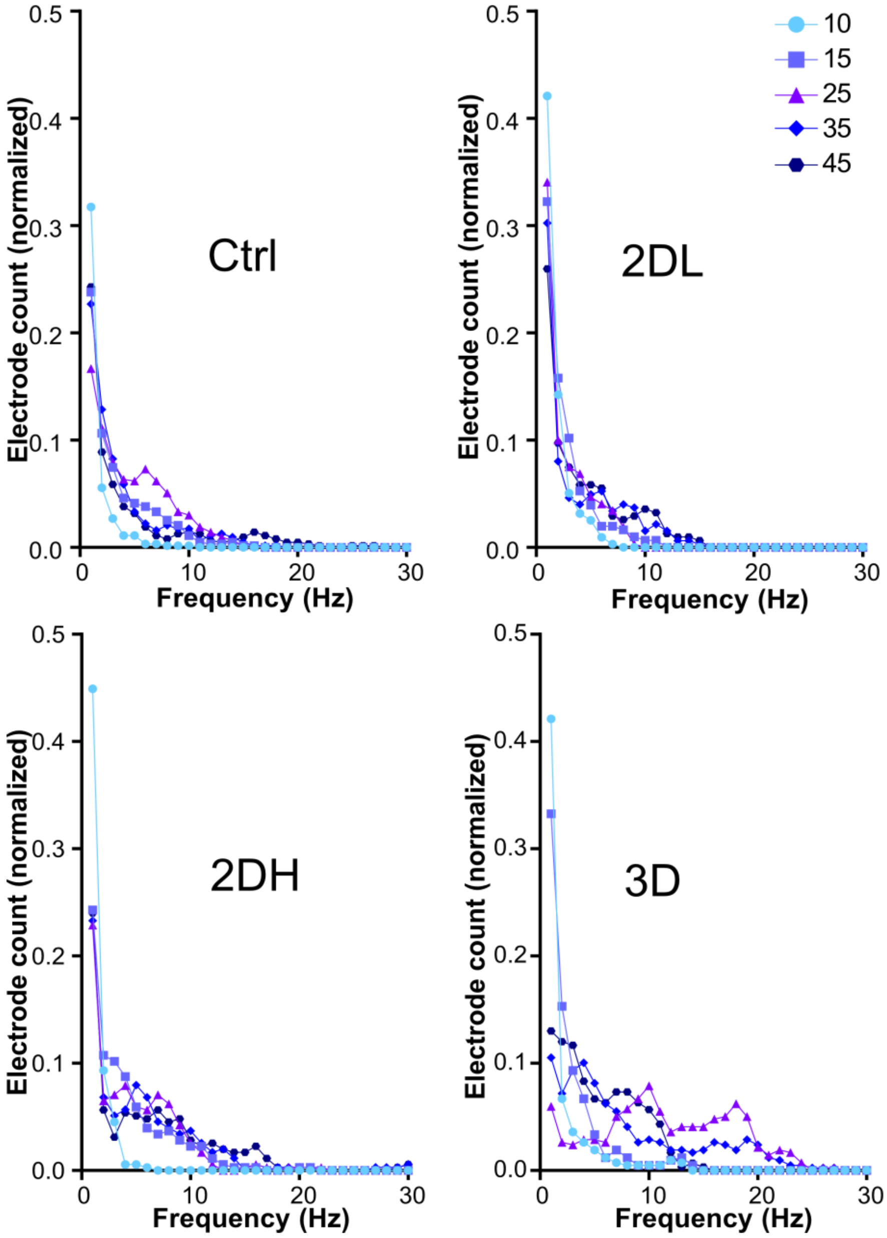


**Supp Fig. 12 Distribution of number of electrodes based on their AP frequencies in different days**. AP frequencies detected by different electrodes were between 0 and 30 Hz and were categorized into 30 bins of 1Hz. The number of electrodes with same AP frequency domain was counted and normalized to the total number of electrodes. For each DIV data have been pooled from 4 MEAs and 236 electrodes per group.

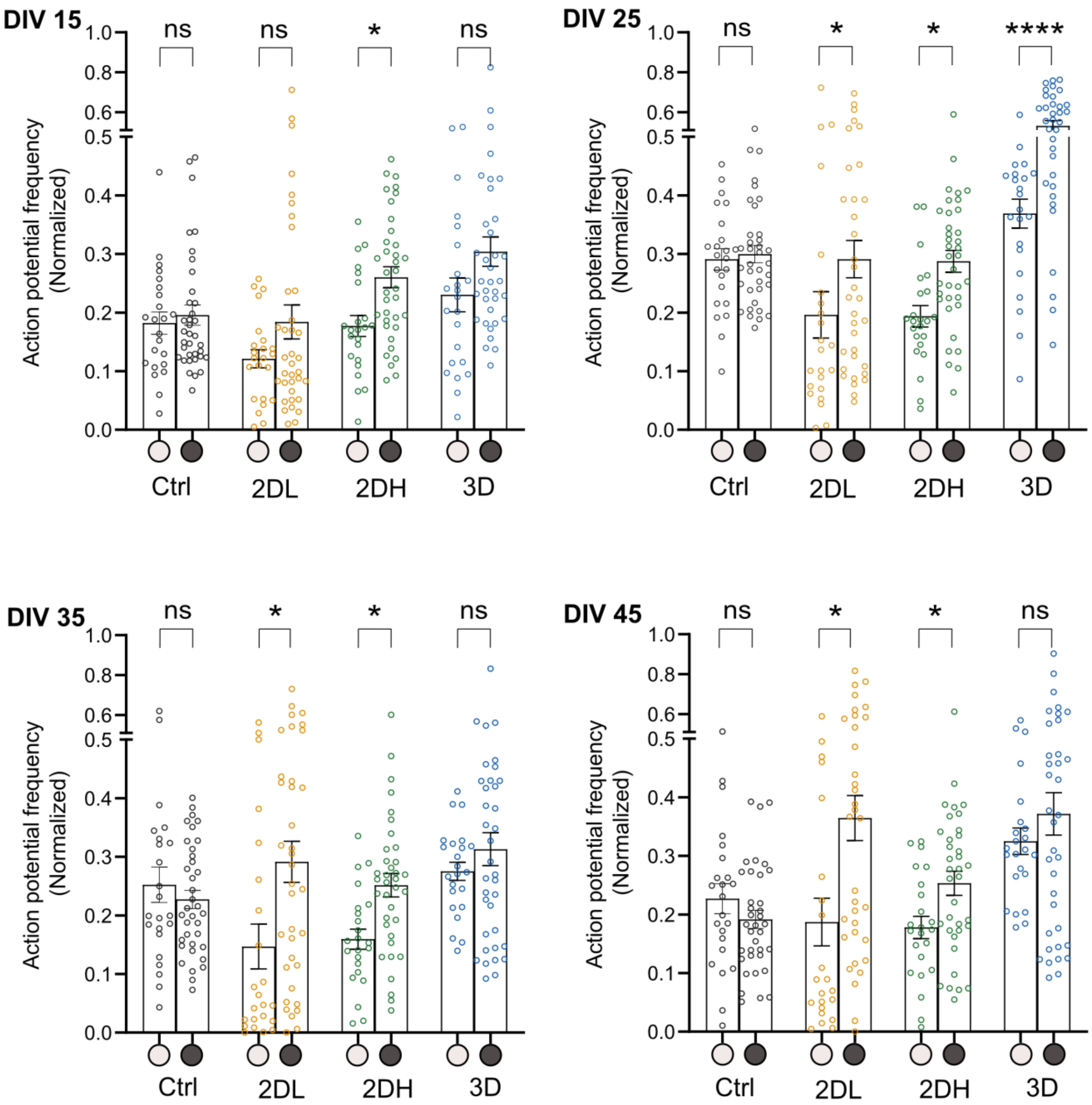


**Supp Fig. 13 Normalized action potential (AP) frequency in 3-edge vs. 4-edge nodes of the same group at different Divs.** The average AP frequency in each node was normalized to the maximum values in the same MEA at that DIV. Data compared between 3-edge vs. 4-edge nodes of the same group using paired t-test (* $p < 0.05$ and **** $p < 0.0001$).

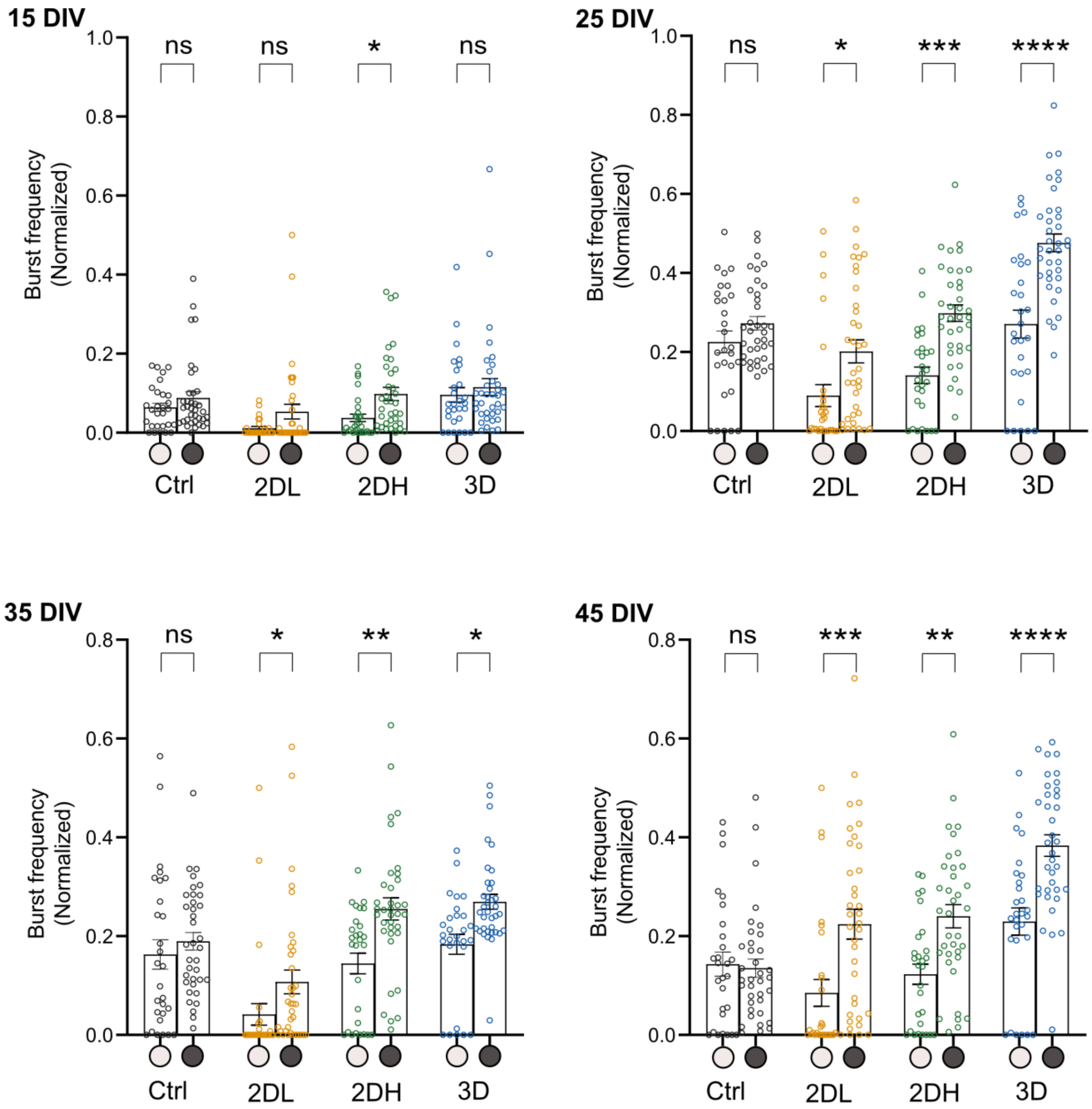


**Supp Fig. 14 Normalized burst frequency in 3-edge vs. 4-edge nodes of the same group at different DIVs.** The average burst frequency in each node was normalized to the maximum values in the same MEA at that DIV. Data compared between 3-edge vs. 4-edge nodes of the same group using paired t-test (* $p < 0.05$, ** $p < 0.01$, *** $p < 0.001$, and **** $p < 0.0001$).

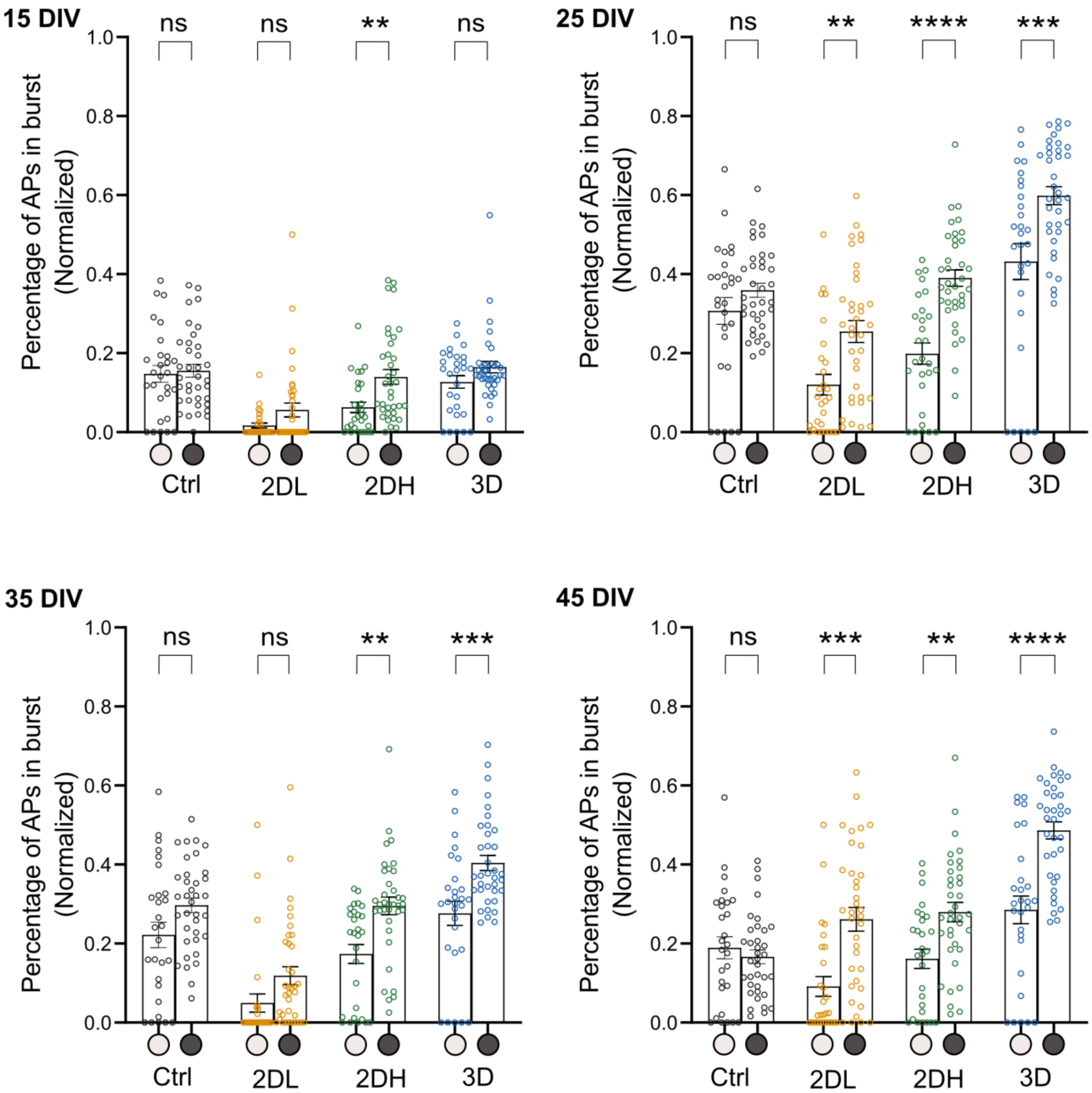


**Supp Fig. 15 Normalized percentage of APs in a burst in 3-edge vs. 4-edge nodes of the same group at different DIVs.** The average of the percentage of APs in a burst in each node was normalized to the maximum values in the same MEA at that DIV. Data compared between 3-edge vs. 4-edge nodes of the same group using paired t-test (** $p < 0.01$, *** $p < 0.001$, and **** $p < 0.0001$).

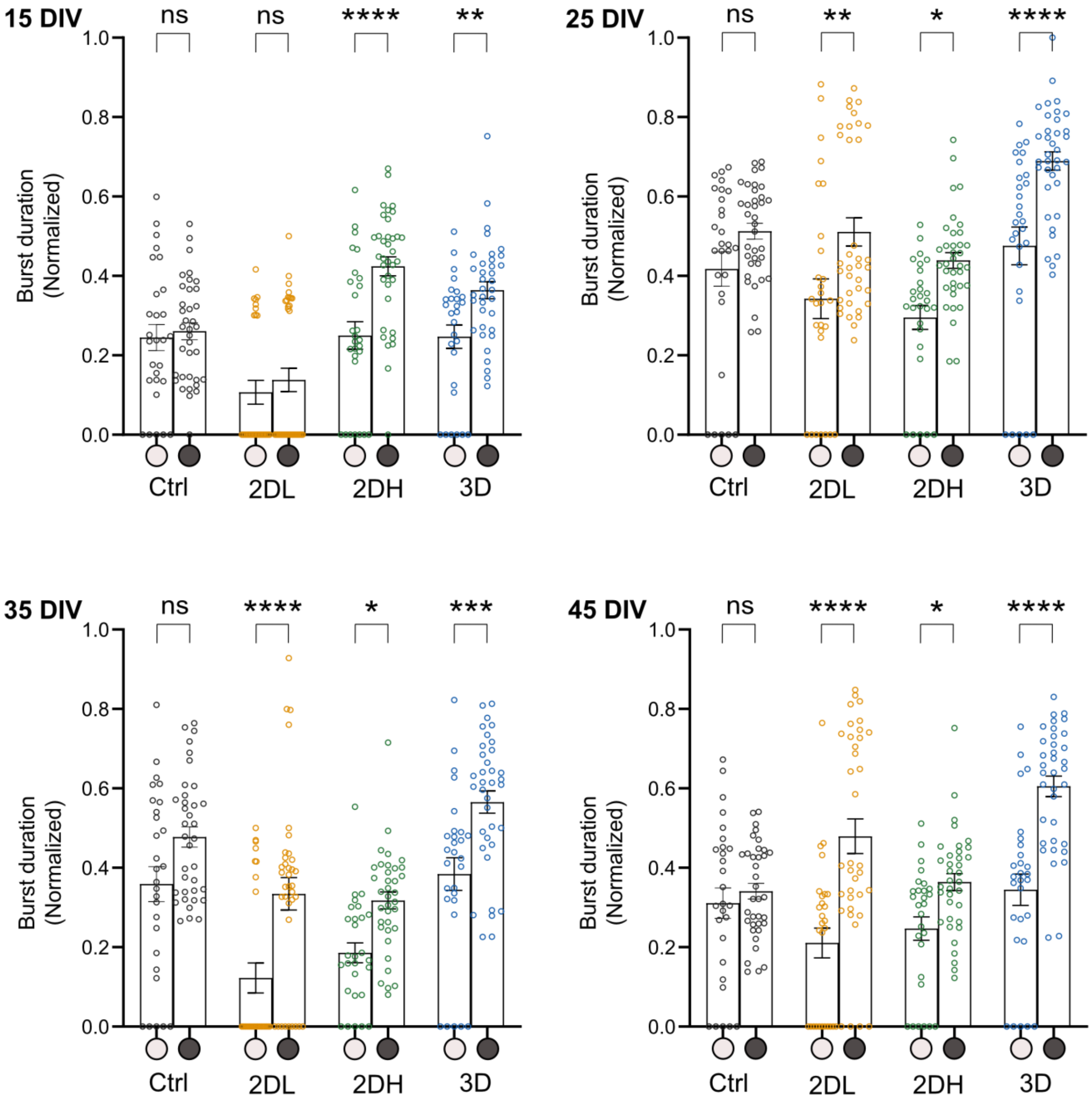


**Supp Fig. 16 Normalized burst duration in 3-edge vs. 4-edge nodes of the same group at different DIVs.** The average burst duration in each node was normalized to the maximum values in the same MEA at that DIV. Data compared between 3-edge vs. 4-edge nodes of the same group using paired t-test (* $p < 0.05$, ** $p < 0.01$, *** $p < 0.001$, and **** $p < 0.0001$).

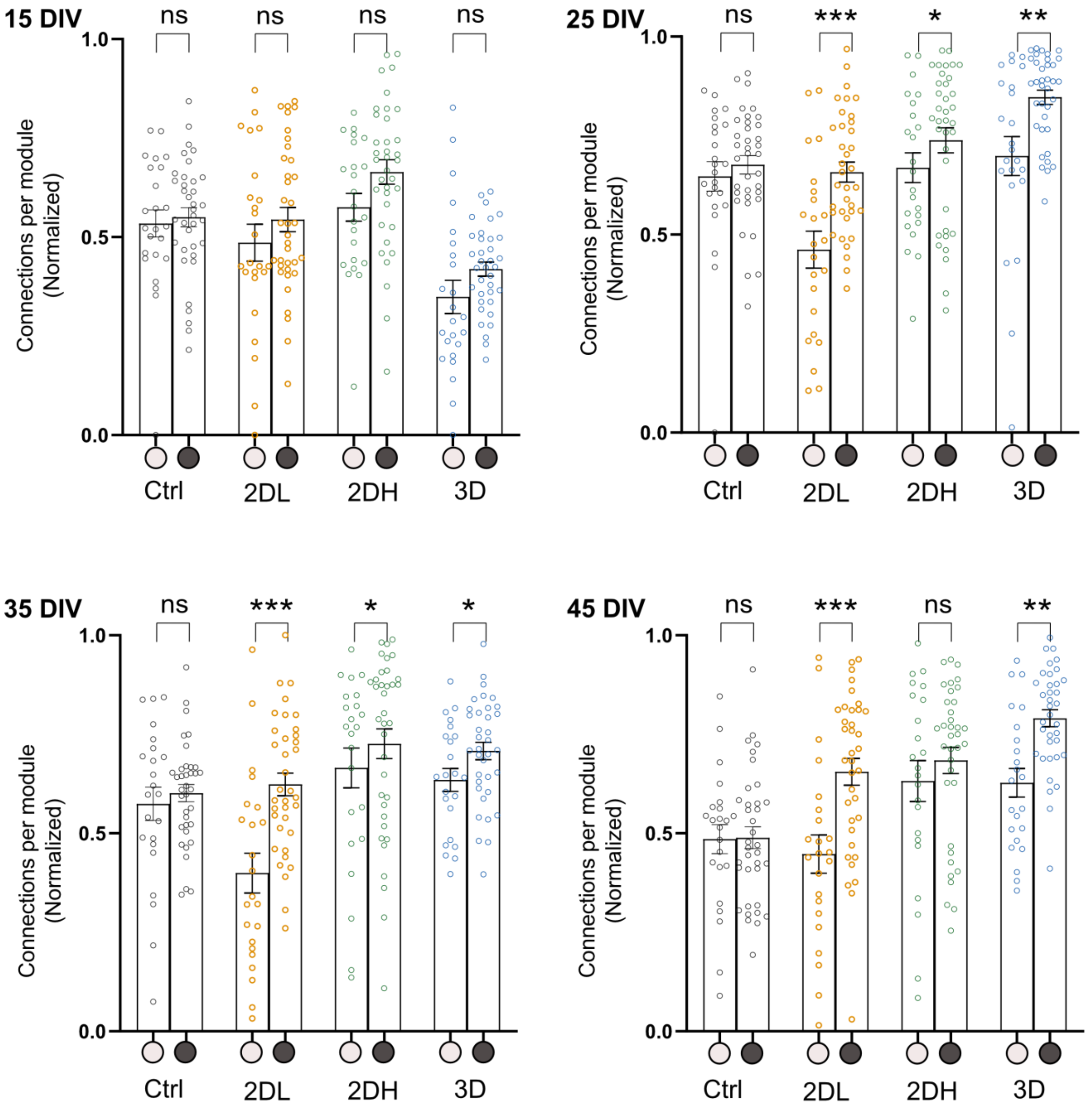


**Supp Fig. 17 Normalized number of connections per module in 3-edge vs. 4-edge nodes of the same group at different DIVs.** The average number of connections per module was normalized to the maximum values in the same MEA at that DIV. Data compared between 3-edge vs. 4-edge nodes of the same group using paired t-test (* $p < 0.05$, ** $p < 0.01$, *** $p < 0.001$, and **** $p < 0.0001$).

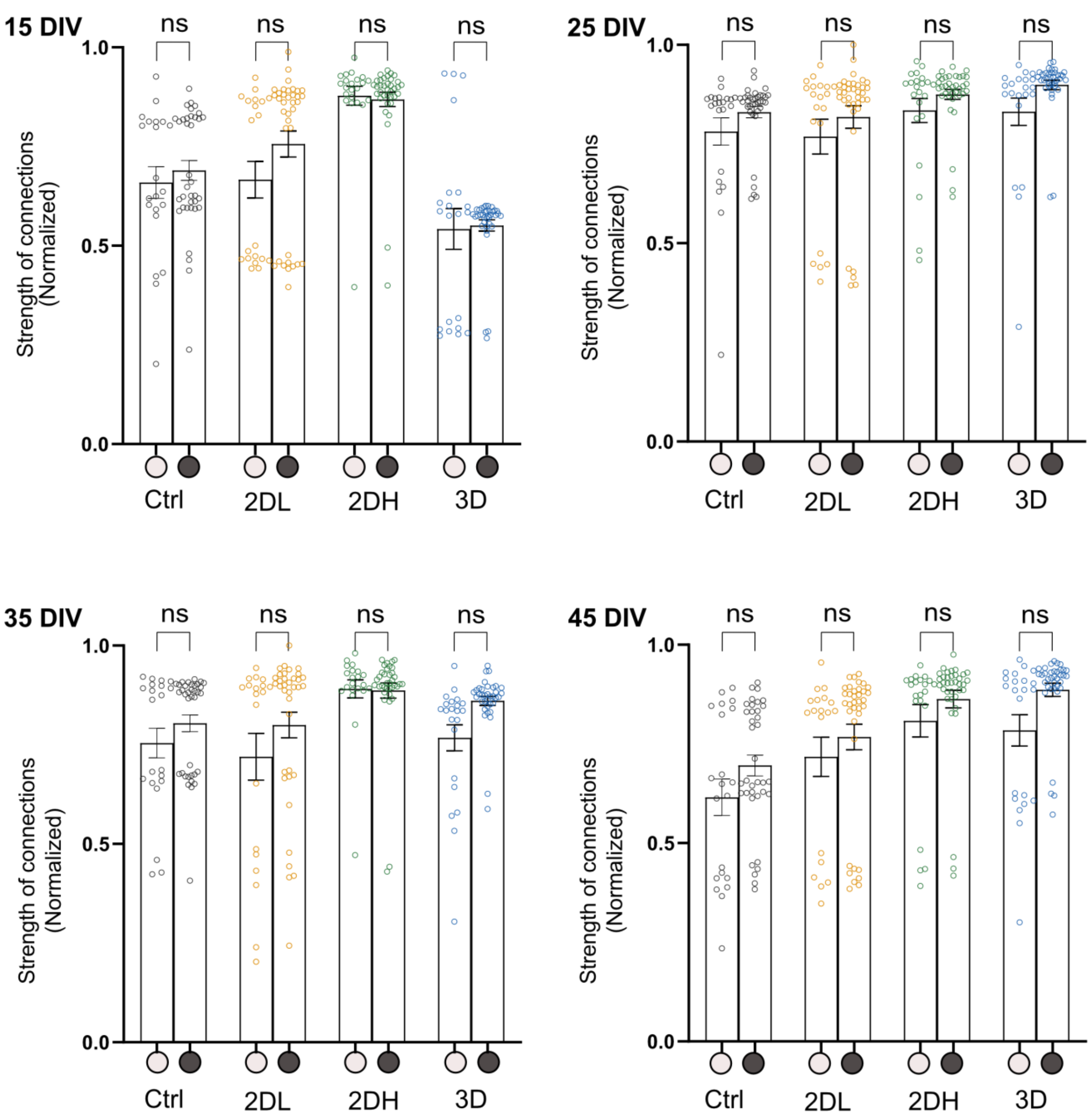


**Supp Fig. 18 Normalized strength of connections per module in 3-edge vs. 4-edge nodes of the same group at different DIVs.** The average strength of connections per module was normalized to the maximum values in the same MEA at that DIV. Data compared between 3-edge vs. 4-edge nodes of the same group using paired t-test (* $p < 0.05$, ** $p < 0.01$, *** $p < 0.001$, and **** $p < 0.0001$).

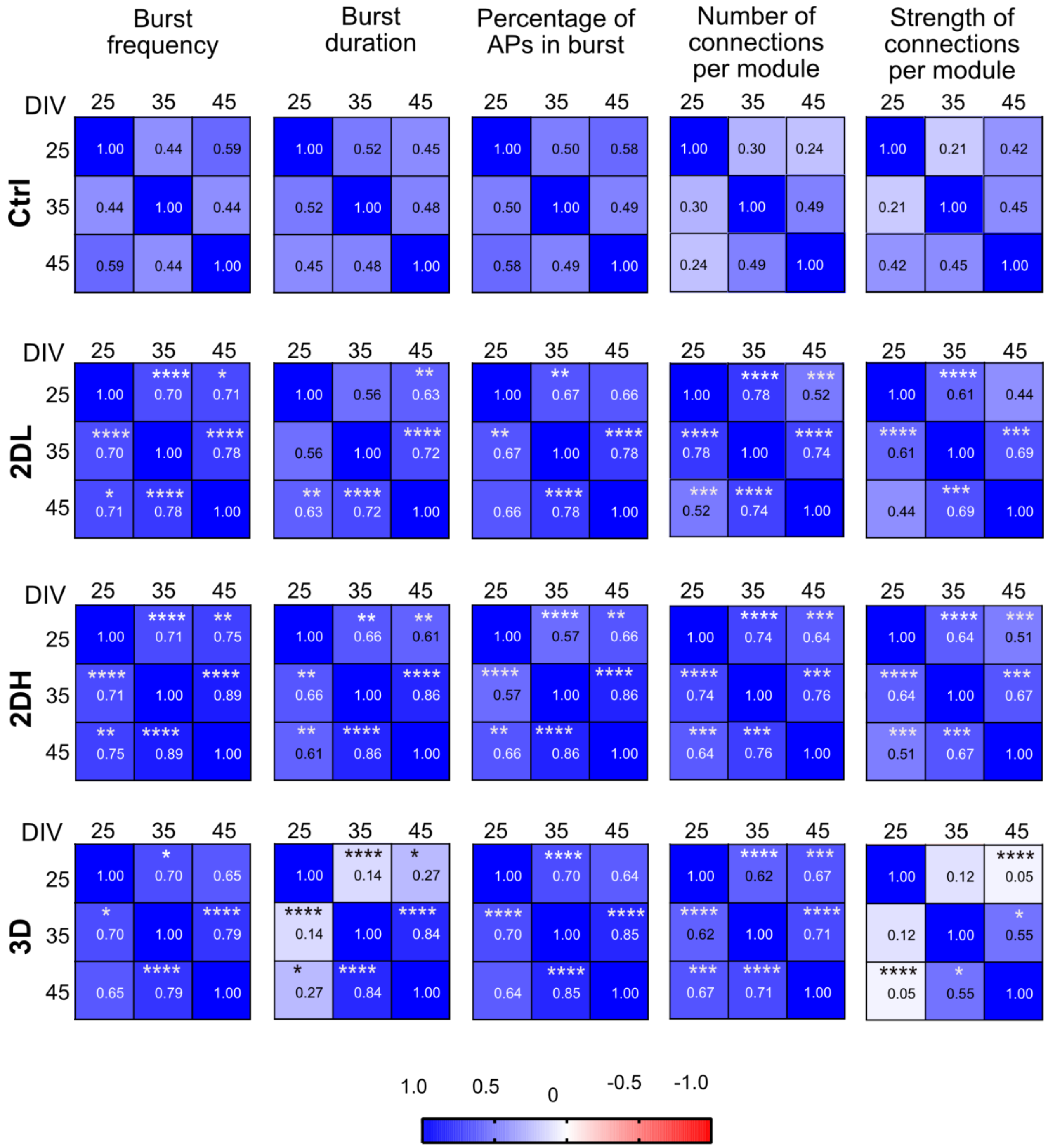


**Supp Fig. 19 Correlation of network parameters over time.** Correlation (Spearman's correlation) between normalized values of different burst features and functional connectivity parameters in the same electrode at different days over network development (N = 4 MEAs per group and n = 236 electrodes per group). Heatmaps and numbers in each graph represent Spearman's r-value (correlation coefficient) between days. Fisher r to z transformation followed by student t-test was applied to compare the 2D or 3D patterned networks vs. random networks. * $p < 0.05$, ** $p < 0.01$, *** $p < 0.001$, and **** $p < 0.0001$.
